# SMC modulates ParB engagement in segregation complexes in *Streptomyces*

**DOI:** 10.1101/2024.10.17.618854

**Authors:** Katarzyna Pawlikiewicz, Agnieszka Strzałka, Michał Majkowski, Julia Duława-Kobeluszczyk, Marcin Szafran, Dagmara Jakimowicz

## Abstract

ParB is long established chromosome segregation protein in bacteria. Due to the recently demonstrated CTPase activity of ParB, formation of its nucleoprotein complexes was unrevealed. ParB homodimers bound to CTP are loaded onto DNA at *parS* sites, where they recruit condensin (SMC), thereby facilitating chromosome organization and segregation. Whether SMC modulates ParB complexes has remained unknown. Here, we generated *Streptomyces venezuelae* strains producing ParB-HaloTag in the presence or absence of SMC and used single-cell time-lapse fluorescence microscopy, single molecule tracking and fluorescence recovery after photobleaching analysis to explore ParB dynamics. Additionally, we performed chromatin immunoprecipitation to examine ParB interactions with DNA in the presence or absence of SMC. We reveal that SMC modulates ParB complex stability on DNA. We find that the absence of SMC results in faster ParB complex disassembly, and promotes non-specific DNA binding. Additionally, we show that SMC reduces ParB CTPase activity *in vitro*. Taken together our data provide evidence of SMC positive feedback on the ParB nucleoprotein complex, offering new insight into the nature of ParB complex regulation.

## INTRODUCTION

The molecular mechanism of bacterial chromosome segregation is not fully understood. In numerous bacterial species, chromosome segregation is driven by the ParABS system (Jalal & Le, 2020; Kawalek et al., 2020; Pióro & Jakimowicz, 2020). ParB forms the nucleoprotein segregation complex (segrosome) by binding *parS* sites that are clustered in the proximity of the origin of chromosome replication (*oriC*) (Breier & Grossman, 2007; Graham et al., 2014; Ireton K et al., 1994; Jalal & Le, 2020). ParB complexes recruit the structural maintenance of chromosomes (SMC) protein (Sullivan et al., 2009; Wang et al., 2015, 2017). SMC is a loop extrusion motor that organizes chromosomal DNA (Ganji et al., 2018; Marko et al., 2019; Ryu et al., 2022; Wang et al., 2015). Moreover, ParB complexes interact with ParA, ATPase that assists their segregation (Jalal & Le, 2020; Kawalek et al., 2020; Lutkenhaus, 2012; Pióro et al., 2022).

Unexpectedly, ParB was recently demonstrated to be a CTPase, opening new avenues for understanding segrosome assembly (Osorio-Valeriano et al., 2019; Soh et al., 2019). According to current models, ParB binds *parS* sites as a CTP-bound homodimer (referred to as the “nucleation stage”) and subsequently undergoes a conformational change to form a closed clamp structure (Osorio-Valeriano et al., 2019, 2021). In this clamp conformation, ParB slides along the DNA, spreading away from the *parS* sites, which drives higher-order DNA-ParB complexes (Jalal et al., 2021). CTP hydrolysis then releases ParB from the DNA, which is accompanied by a conformational change. CTP hydrolysis deficient ParB variants exhibited excessive spreading indicating that nucleotide hydrolysis increases the turnover of protein (Antar et al., 2021). While the role of CTP binding and hydrolysis was established, some of the details of ParB complex formation on DNA are still elusive. For example, the mechanism of ParB binding to regions flanking several kilobases from *parS* sites is still under debate (Balaguer et al., 2021; Connolley et al., 2023; Tišma et al., 2024). Furthermore, ParB ability to undergo liquid-liquid phase separation and form condensates was brought up to contribute to segrosome assembly (Babl et al., 2022; Zhao et al., 2024). Finally, the impact of SMC loading on the ParB complex and its interaction with ParA remains unexplored.

The ParB complex was demonstrated to recruit the SMC protein in numerous bacterial species (Gruber & Errington, 2009; Minnen et al., 2011; Sullivan et al., 2009; Wang et al., 2015). In *Bacillus subtilis*, SMC was reported to directly interact with the N-terminal domain of ParB, which is also the domain engaged in ParA and CTP binding (Bock et al., 2022). Interestingly, SMC recruitment by ParB depends on its CTP binding but does not require CTP hydrolysis (Antar et al., 2021). The interactions between SMC and ParB are proposed to affect ParB association with ParA (Antar et al., 2021; Taylor et al., 2021). By interaction with ATP-bound ParA dimer associated nonspecifically with DNA, the ParB complex stimulates ParA ATPase activity and releases ParA from the nucleoid. This mechanism provides the driving force for chromosome segregation. The contribution of ParA to SMC loading was shown, indicating the crosstalk between both ParB binding partners, ParA and SMC (Roberts et al., 2024).

In *Streptomyces*, a multigenomic, filamentous and sporulating bacteria, the ParB and SMC proteins organize multiple chromosomal copies in a developmental stage-dependent manner (Szafran et al., 2020, 2021). ParB binds to numerous (16 in *S. venezuelae*) *parS* sites located near the *oriC* region at the centre of the linear *Streptomyces* chromosome (Donczew et al., 2016; Jakimowicz et al., 2002). In apically growing and branching vegetative cells, the main role of ParB complexes is to anchor *oriC* of the apical chromosome to the cell pole (hyphal tip), thereby facilitating efficient tip extension and branching. This anchorage of tip-associated ParB complex requires ParA to interact with polar proteins (polarisome, also known as a tip-organizing complex, TIPOC) (Ditkowski et al., 2013; Hempel et al., 2008; Holmes et al., 2013; Kois-Ostrowska et al., 2016). During sporulation, which turns elongated sporogenic cells into chains of unigenomic exospores, multiple chromosomal copies are segregated and compacted. The extensive elongation of the sporogenic cell accompanied by intensive chromosome replication is followed by growth arrest. Next, the sporogenic cell undergoes multiple synchronized cell divisions. At this stage, ParA and ParB ensure efficient segregation of chromosomes into spores (Donczew et al., 2016; Jakimowicz et al., 2007; Kim et al., 2000). ParA accumulates along sporogenic cells while ParB forms an array of uniformly distributed segrosomes. Recently, it was shown that in *Streptomyces*, as in other bacteria, ParB complexes recruit SMC (Szafran et al., 2021). The SMC recruitment is required for efficient chromosome compaction within the spores. SMC loading by ParB rearranges *Streptomyces* chromosomes during sporogenic development from an open conformation with limited interarm contacts to a closed conformation with both arms aligned in spores (Szafran et al., 2021).

Given that ParB complexes in *Streptomyces* recruit SMC, we wondered if SMC loading would impact the positioning and dynamics of ParB complexes. We expected that the loading of SMC which is enhanced in sporogenic cells would facilitate the progress of chromosome segregation. Therefore, we labelled ParB with HaloTag (ParB-HT) in *S. venezuelae* strains, taking advantage of the model species that allows the application of the time-lapse microscopy for analysis of sporogenic development. We followed the dynamics of ParB-HT complexes in sporogenic hyphae of *S. venezuelae* in the presence and absence of SMC. We found that the absence of SMC shortens the lifetime of ParB-HT segregation complexes but also reduces the mobility of ParB-HT. ParB-DNA binding studies in wild type and *smc* mutant strains show that SMC promotes ParB binding to *parS* and spreading, while measurements of ParB CTPase activity demonstrate that SMC reduces CTP hydrolysis by ParB. Together, our findings reveal previously unknown features of ParB complexes, establishing that SMC loading has positive feedback on segrosome stability.

## RESULTS

### The dynamics of ParB complexes in sporogenic cells

To study the dynamics of ParB complexes during sporogenic chromosome segregation of *S. venezuelae*, we labelled ParB by fusing it with HaloTag (HT). The levels of ParB-HT were similar to the levels of wild-type ParB, either for ParB-HT produced from a repressible promoter or from the native *parAB* promoter (Δ*parB*, p_rtet_*parB-HT*, strain KP009, or Δ*parAB*, p_nat_*parAB-HT*, strain KP006, respectively, for details see the Supplementary Information). The ParB-HT fusion protein was confirmed to be functional and had no adverse effects on the growth of the KP009 or KP006 strains **(Fig. S1A and B)**. Fluorescence microscopy detecting Halo-Tag labelling with TMR ligand revealed the presence of irregularly spaced ParB-HT complexes in vegetative hyphal cells, with a characteristic bright focus at the hyphal tip representing apical ParB complex **(Fig. S1C)**, consistent with previous analyses of *S. coelicolor* (Kois-Ostrowska et al., 2016). To determine the dynamics of ParB complexes during sporogenic development, the ParB-HT strain (KP009) was further modified to produce cell division protein FtsZ fused with YPet (strain KP011) to mark Z-ring formation during cytokinesis, as described earlier (Donczew et al., 2016; Schlimpert et al., 2017).

The ParB-HT complex dynamics in sporogenic cells of the KP011 strain was followed with time-lapse single-cell fluorescent microscopy. ParB-HT was visible as irregularly spaced complexes during sporogenic cell extension (**Fig. 1A, B, Fig. S2, Fig. S3)**. Shortly after cell extension stopped (T1 = ∼27 +/− 10 min), these distinct ParB-HT complexes became regularly spaced along the hyphal cell (resembling those observed earlier in *S. coelicolor* (Jakimowicz et al., 2005) **(Fig. 1A, Fig. S2, Fig S3)**. Noticeably, for the next hour, the ParB complexes gradually diffused and fully disassembled approximately 106 +/− 16 min after cell growth arrest (T2). Thus, regularly spaced segrosomes lasted approximately 80 min. The regularly spaced ParB-HT complexes accompanied Z-rings, which also formed approximately 27 +/− 12 min (T3) after cell growth arrest and lasted approximately 80 min +/− 10 **(Fig. 1A-C)**. Thus, ParB-HT complexes disassemble at the same time as Z-rings disassemble during *S. venezuelae* sporogenic cell division.

**Figure 1.**
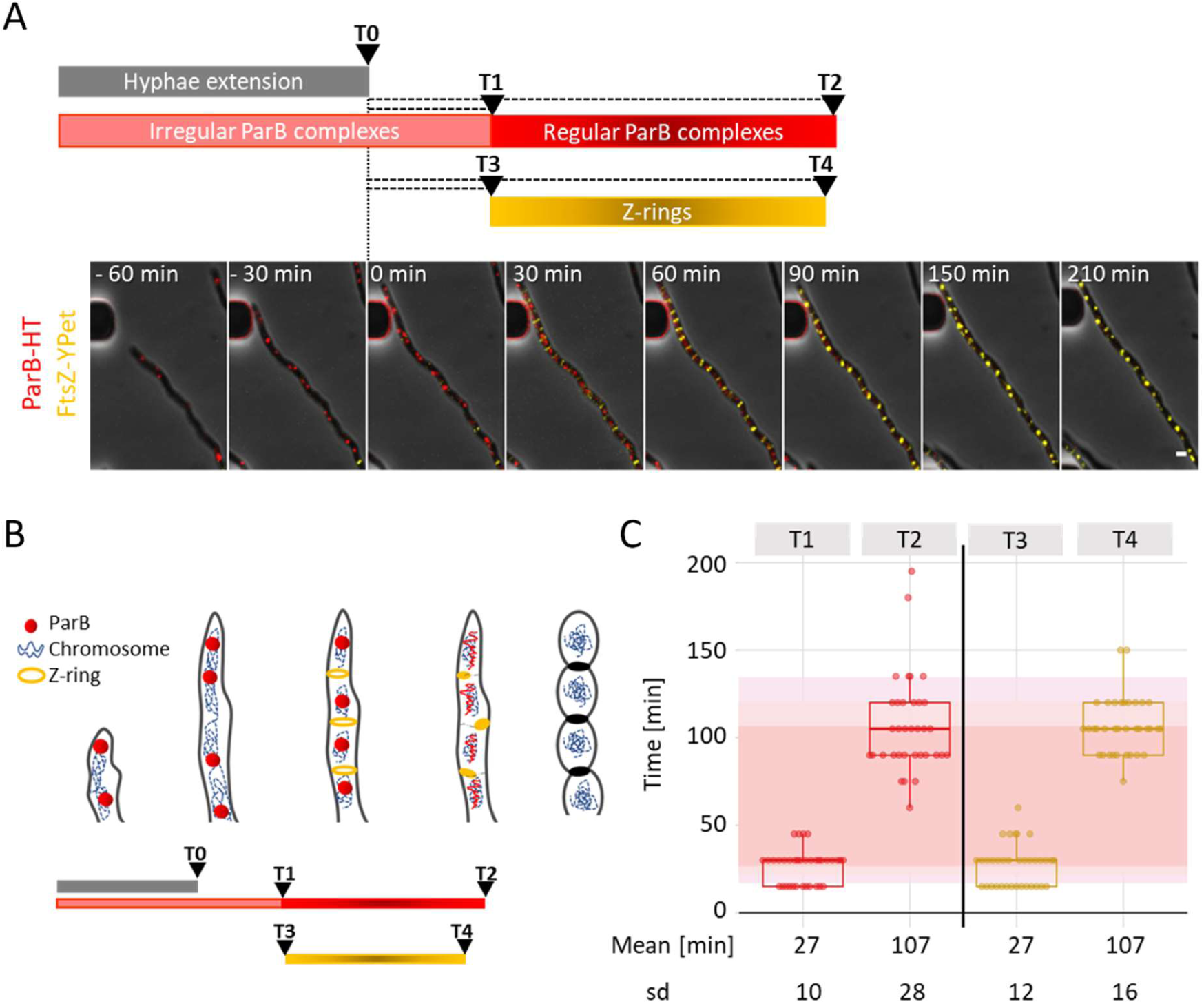
ParB-HT complexes in *S. venezuelae* sporogenic cells are short-lived and disassemble during cell division. **A**. Representative images from time-lapse analysis of sporogenic development of Δ*parB* p_rtet_*parB-HT ftsZ-ypet* (KP011) strain showing fluorescence of ParB-HT stained with Janelia Fluor-549 (red) overlaid with FtsZ-YPet fluorescence (yellow) and phase contrast (grey). In A and B time 0 is the time of sporogenic cell growth arrest and the analysed stages of sporulation are indicated in the bar scheme below. Scale bar – 1 μm. For separate channels see Fig. S2. **B.** Scheme of the sporogenic cell development indicating the stages which time was measured **C**. Analyses of the time elapsed from growth cessation (time 0) to appearance of regularly spaced ParB-HT complexes (T1), their disappearance (T2), appearance of regularly spaced Z-rings (T3) and their disappearance (T4). Red shading shows the mean lifetime of ParB complexes. Data shown in panel C were collected in 3 independent experiments for 35 hyphae.

### Elimination of SMC promotes disassembly of ParB-HT complexes in sporogenic cells

We investigated whether SMC loading could affect segrosome dynamics. To test this, we compared the sporogenic development of the strain producing ParB-HT in the Δ*smc* genetic background (Δ*smc*Δ*parAB*p_nat_*parAB-HT*, strain KP007) with the wild type control (Δ*parAB*p_nat_*parAB-HT*, KP006 strain) using single-cell time-lapse microscopy. Of note, ParB-HT (produced from p_nat_*parAB-HT*) complemented severe chromosome segregation defects, detected in τηε Δ*smc*Δ*parAB* strain **(Fig. S1D).**

In sporogenic cells lacking *SMC* (KP007 strain), regularly spaced ParB-HT complexes were detected marginally earlier (T1 = 20 +/− 9 min from hyphae growth cessation) than in the wild type control (KP006 strain) (T1 = 29 +/− 9 min) **(Fig. 2A-C, Fig. S4)**. Strikingly, the ParB complexes disassembled significantly earlier in the absence of SMC (T2 = 79 +/− 25 min from hyphae growth cessation), than in the control strain (T2 = 114 +/− 42 min) **(Fig. 2, Fig. S4)**. Notably, in the absence of SMC, the lifetime of ParB complexes was not only shortened but also showed lower variation than in the wild type background, while the distances between the segrosomes were not affected (**Fig. S3**).

**Figure 2.**
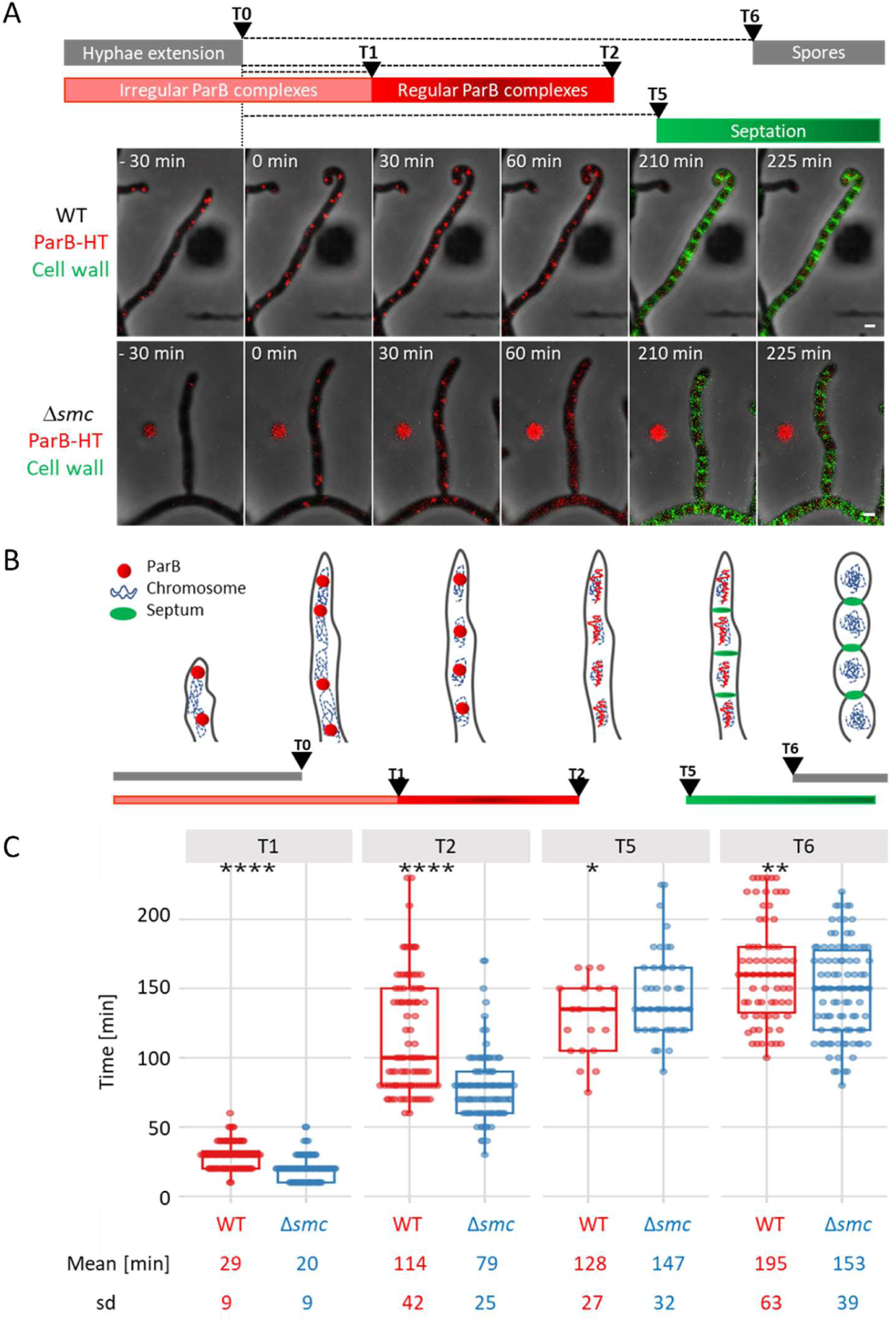
The elimination of SMC accelerates disassembly of ParB-HT complexes in *S. venezuelae* sporogenic hyphae. **A**. Representative images from time-lapse analysis of sporogenic development of Δ*parAB* p_nat_*parAB-HT*, KP006 strain and Δ*smc*Δ*parAB* with p_nat_*parAB-HT*, KP007 showing fluorescence of ParB-HT stained with Janelia Fluor-549 (red) overlaid with fluorescence of NADA green stained septa (green) and phase contrast (grey). In A and B time 0 is the time of hyphal cell growth arrest. Scale bar – 1 μm. **B**. Scheme of analysed time intervals. **C**. Comparison of the time elapsed from growth cessation (time 0) to appearance of regularly spaced ParB-HT complexes (T1), their disappearance (T2), the appearance of septa (NADA signal) (T5) and spores (T6) in the wild type control and Δ*smc* background (KP006 and KP007). Data shown in panel D were collected in 3, or 2 for NADA stained hyphae, independent experiments for 96 hyphae of KP006 strain and 99 hyphae o KP007 strain (to determine T1, T2, T6) and 19 hyphae of KP006 strain and 43 hyphae of KP007 strain (to determine T5), statistical analyses were performed using a two-sided t-student test.

To determine the time of sporogenic cell division, ParB-HT producing strains were labelled with fluorescent D-amino acid (NBD-amino-d-alanine, NADA) in the time-lapse microscopy analyses. In the wild type control strain (KP006), septation (NADA signal) was detected approximately 128 +/− 27 min (T5) after hyphal growth cessation and 14 (+/− 32) minutes after ParB complex disassembly. The absence of SMC slightly delayed the time of septation (T5 = 147 +/− 32 min after cell growth cessation) compared with the control strain **(Fig 2A-C)**. This was consistent with delayed Z-ring disassembly observed in τηε Δ*smc* background **(Fig. S5)**. Despite the delayed cell division, the overall time of sporulation (from growth arrest to detection of rounded spores, T6) was only marginally affected by the absence of SMC **(Fig. 2, Fig. S4)**.

The shortened lifetime of ParB-HT complexes in the absence of SMC could result from the lowering of ParB-HT levels. However, Western blotting of ParB-HT levels in KP006 and KP007 strains during their sporogenic development showed that they were close to constant during sporogenic development and that they were not affected by the elimination of SMC **(Fig. S6)**. Thus, changes in ParB-HT levels cannot account for the rapid disassembly of ParB complexes in the absence of SMC. Overall, these findings indicate that SMC stabilizes the ParB complex during sporogenic development.

### Elimination of SMC lowers mobility of ParB molecules in sporogenic and vegetative cells

Next, we utilized single-molecule tracking to determine how *smc* deletion affects ParB-HT mobility. Earlier studies based on tracking various DNA binding proteins demonstrated that increased DNA association often results in reduced mobility (Garza de Leon et al., 2017; Kapanidis et al., 2018). Accordingly, we expected that the disassembly of ParB-HT complexes would be reflected by increased mobility. ParB-HT mobility was studied at different stages of development: early sporogenic cells (17 h of liquid culture, where most of the sporogenic hyphae contained ParB-HT complexes), late sporogenic cells (22 h of liquid culture, where in some of the hyphae ParB-HT complexes disassembled) **(Fig. S7)**, as well as in spores. We found that single-step distances (the distance that single molecules cover between two time points) raised during differentiation **(Fig. 3A).** Similarly, the average apparent diffusion coefficient (d) increased from 0.052 μm^2^/s in the early stage of sporulation to 0.106 μm^2^/s in the late sporogenic cell (**Fig. 3B**). ParB-HT mobility was the highest in spores (d = 0.220 μm^2^/s) **(Fig 3A, B)**. These changes in ParB-HT mobility are consistent with the observed segrosome disassembly.

**Figure 3.**
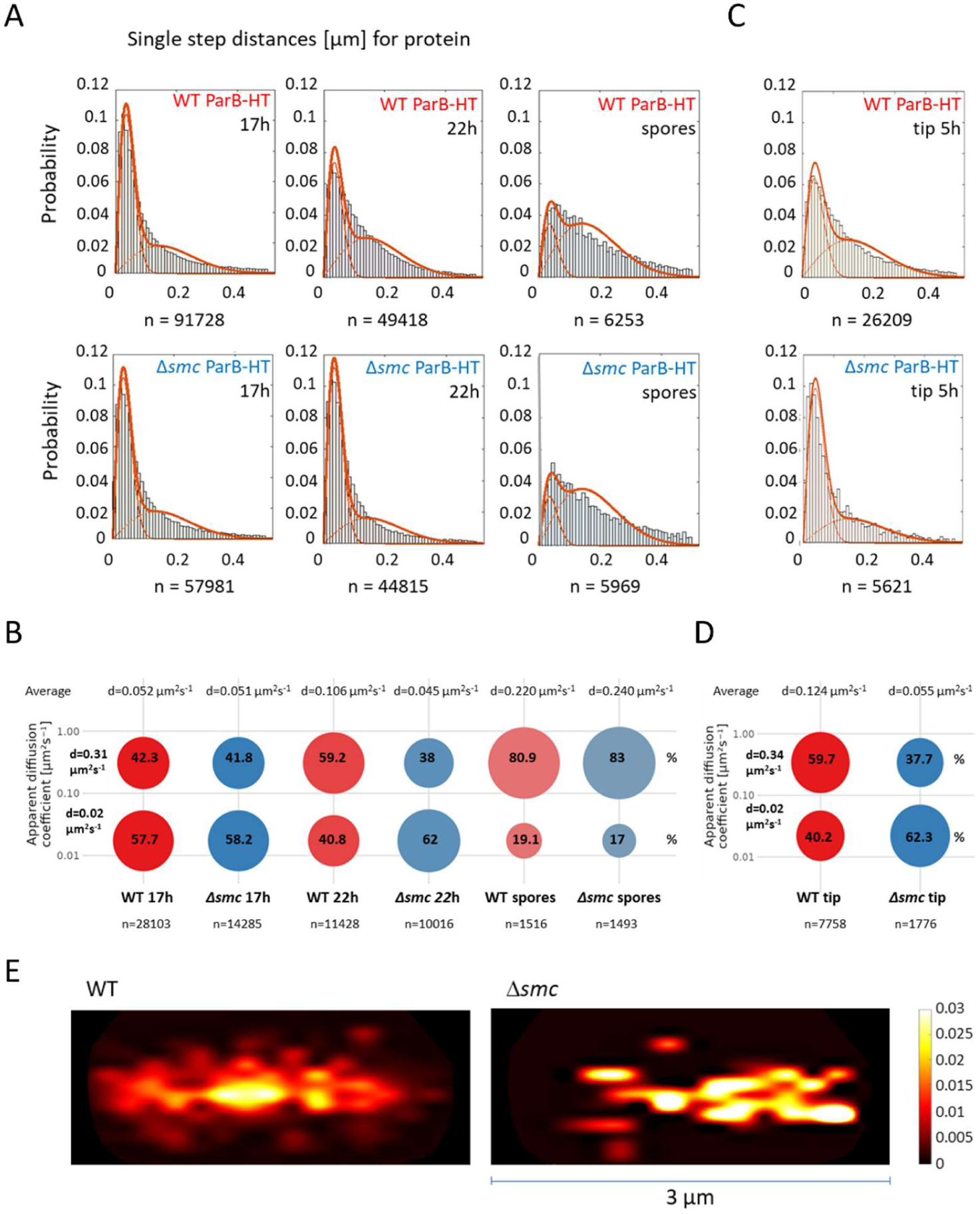
Mobility of ParB-HT is lowered by the absence of SMC. **A.** single step distance of ParB-HT in sporogenic cell of the wild type control (KP006) and Δ*smc* strains (KP007) at different stages of sporogenic development **B.** Percentage of ParB-HT tracks with low (d=0.02 μm^2^/s) and high (d=0.31 μm^2^/s) diffusion coefficient at different stages of sporogenic development. **C.** Single step distance of ParB -HT analysis in young vegetative cells (3 µm distance from the tip) of the wild type control (KP006) and Δ*smc* strains (KP007) **D.** Percentage of ParB-HT tracks with low (d=0.02 μm^2^/s) and high (d=0.34 μm^2^/s) diffusion coefficient in young vegetative cells (3 µm distance from the tip) of the wild type control (KP006) and Δ*smc* (KP007) strains. **E**. Heatmap showing a number of tracks with apparent diffusion MSD between 0-0.13 µm^2^ in young vegetative cells of the wild type control and Δ*smc* strains. Data shown in A-D were collected in 2 independent experiments for wild type control and Δ*smc* strains (Δ*parAB* p_nat_*parAB-HT*, KP006 and Δ*smc*Δ*parAB* p_nat_*parAB-HT* KP007, respectively). A, C - Two population model fit and number of analysed steps are shown on each histogram, B, D - Average apparent diffusion coefficient (d) and number of analysed tracks (n) are shown for each strain.

In early sporogenic cells, *smc* deletion did not change the mobility of ParB-HT molecules (average diffusion coefficient 0.051 μm^2^/s in Δ*smc* background, KP007 vs 0.052 μm^2^/s in wild type control KP006 strain, respectively) **(Fig. 3A, B)**. However, in late sporogenic cells (22 h of growth), the average diffusion coefficient was significantly lower (0.045 μm^2^/s) in the absence of SMC than in the wild type control strain (0.106 μm^2^/s). This was reflected by the increased fraction of ParB-HT molecules with a low diffusion coefficient in Δ*smc* cells (62% of with average d = 0.02 μm^2^/s in Δ*smc* KP007 strain) as compared in wild type control strain (41.5% in KP006) **(Fig. 3A, B)**. Notably, the lack of SMC did not change ParB-HT mobility in spores as compared to wild type control **(Fig. 3A, B)**.

The reduced ParB-HT mobility that could indicate increased DNA binding in late sporogenic cells lacking SMC was surprising because it seemed to contradict the shortened lifetime of segrosomes that we observed in our single-cell time-lapse microscopy analyses. This could be explained by the delayed entry of the Δ*smc* (KP007) strain into sporulation and therefore longer prevalence of hyphae still undergoing sporogenic cell division and chromosome segregation at the 22 h time point. However, at this time point the fraction of Δ*smc* sporogenic cells that showed disassembled ParB-HT complexes was only marginally lower than in the wild type control KP006 strain **(Fig. S7B)**. Moreover, since sporogenic cell division and chromosome segregation follow the shutdown of chromosome replication (Szafran et al., 2021), we performed marker frequency analyses (*ori:arm*) in both strains. This analysis showed no significant difference in chromosome replication rate between the wild type control (KP006)and Δ*smc* (KP007) strains during sporogenic development **(Fig. S7C**). This suggests that minor differences in development did not account for the significant difference in ParB-HT mobility observed in the absence of SMC.

To further explore the impact of SMC on ParB mobility, we tested if *smc* deletion could modulate ParB mobility in early vegetative cells. Western blotting analyses showed that FLAG-SMC levels are constant throughout the life cycle of *S. venezuelae* **(Fig. S8)**. At the early stage of growth (5 hours), hyphal cells contain only a few chromosomes (usually less than 10 ParB foci) and the apical ParB complex is responsible for *oriC* anchorage at the hyphal cell tip. This apical ParB complex remains at a constant distance from the cell pole (∼1.3 μm)(Kois-Ostrowska et al., 2016) allowing for a detailed examination of its dynamics. In early vegetative cells, as in sporogenic cells, the absence of SMC significantly reduced the mobility of ParB-HT (d=0.055 μm^2^/s in Δ*smc* KP007 strain compared to d=0.124 μm^2^/s in the wild type control KP006 strain). Consequently, the fraction of immobile ParB-HT molecules increased (62.3% molecules with average d=0.02 μm^2^/s in Δ*smc* KP007 strain compared to 40.2% in the wild type control KP006 strain) (**Fig. 3C, D, Fig. S9**). Next, we determined the position of immobile molecules (molecular square displacement MSD 0-0.13 µm^2^ in relation to the cell pole. In the wild type control strain, the immobile ParB-HT molecules were constrained to a single location of the cell (∼ 1.5 µm from the hyphal tip), while in the absence of SMC the immobile ParB-HT were more dispersed (**Fig. 3E)**. These results indicate that SMC lowers ParB-HT dynamics at the poles of vegetative cells, but immobile molecules are not engaged in the formation of the single ParB-HT complex.

### The absence of SMC increases the turnover of ParB-HT in the apical complex

We next analysed the dynamics of the apical ParB-HT complex in wild type control and Δ*smc* strains (KP006 and KP007, as above). Since this complex is present throughout the vegetative growth and does not become disassembled, to determine its stability and ParB-HT turnover we used fluorescence recovery after photobleaching (FRAP). While in both analyzed strains (KP006 and KP007) about 60% of ParB-HT complex fluorescence recovered after photobleaching, the time required for half-recovery (tau) was shorter in the absence of SMC (on average 135 s in KP007) than in wild type control strain (on average 222 s in KP006) **(Fig 4A-C, Fig. S10)**. Additionally, in the wild type control strain (KP006), the variation in recovery time was much higher than in the absence of SMC. Shortened recovery time indicates that lack of SMC increases the availability of ParB-HT molecules for complex restoration indicating faster turnover. The observed higher complex turnover rate in the absence of SMC corresponds to the lowered stability of segrosomes in sporogenic cells but contradicts the decreased mobility of ParB-HT determined by single-molecule tracking.

**Figure 4.**
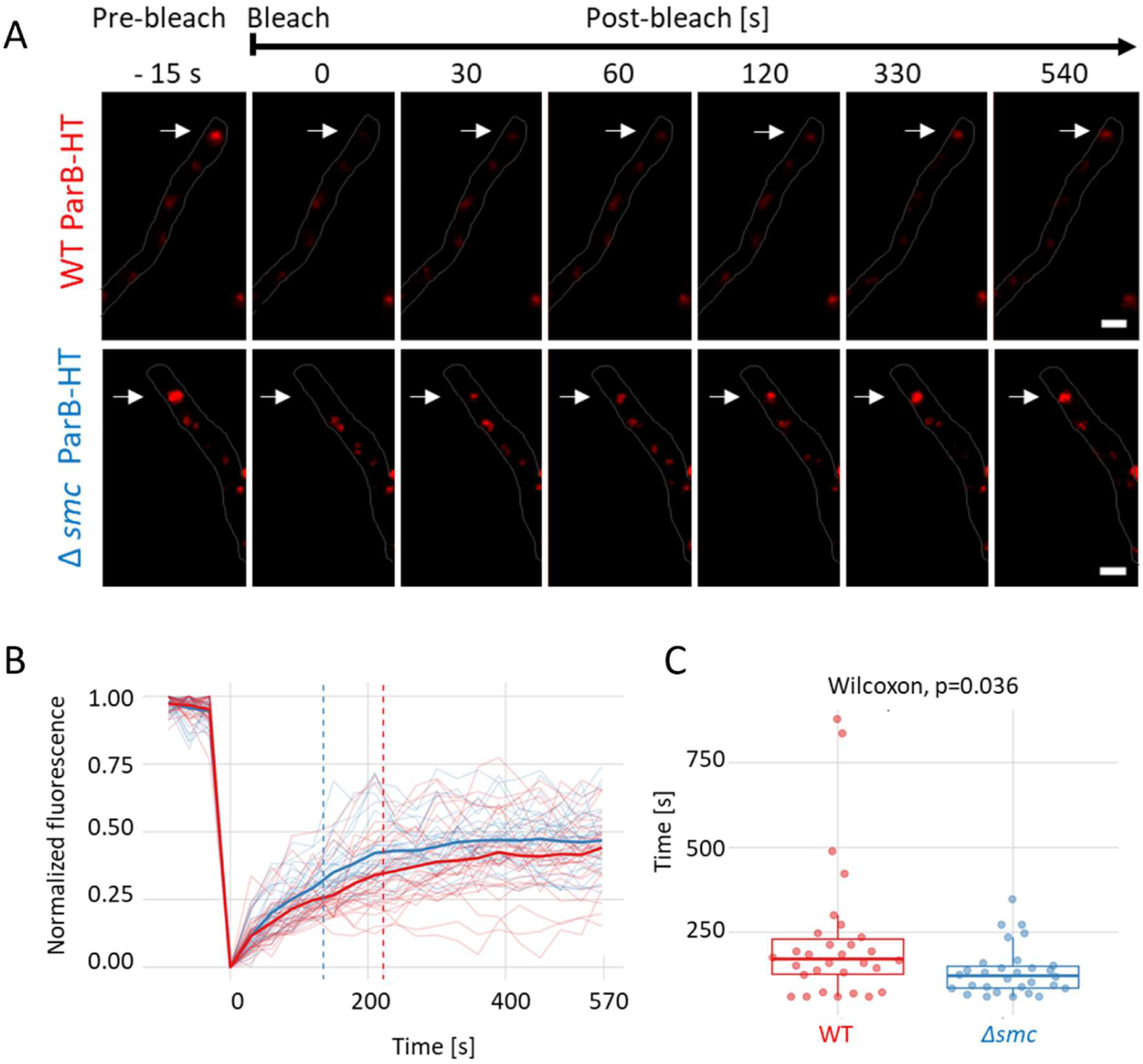
Elimination of SMC shortens the ParB-HT complex recovery time after photobleaching. Representative images showing the photobleaching of the ParB-HT complex stained with Janelia Fluor-549 in young vegetative cells of the wild type control and Δ*smc* strains (Δ*parAB* p_nat_*parAB-HT*, KP006 and Δ*smc*Δ*parAB* p_nat_*parAB-HT* KP007, respectively). ParB-HT fluorescence (red) is overlaid on the brightfield channel (grey) **B.** Fluorescence recovery analysis. Intensity of fluorescence plotted against the time of analyses. The dotted lines show mean Tau values for each strain. **C.** Tau - the time required for recovery of half fluorescence intensity calculated for the wild type control and Δ*smc* strains (Δ*parAB* p_nat_*parAB-HT*, KP006 and Δ*smc*Δ*parAB* p_nat_*parAB-HT* KP007, respectively). Data shown in B and C were collected in 3 independent experiments for 30 ParB-HT complexes of each strain. Statistical analysis was conducted using a two-sided Wilcoxon test.

### SMC promotes ParB spreading at *parS* sites

Having found that the absence of SMC reduces the mobility of ParB-HT but also lowers the stability of ParB-HT complexes, we set out to examine whether the observed differences could be attributed to SMC-mediated modulation of ParB interactions with DNA. To this end, we analysed the binding of ParB (untagged protein) to DNA in the wild type and Δ*smc* strains using ChIP-seq with anti-ParB antibodies (using the Δ*parB* strain, MD020 as a negative control). To eliminate any influence of the potential difference in the sporogenic development stage and because single-molecule tracking showed significantly increased ParB mobility in young vegetative cells lacking SMC (5 hours of growth), our analyses were performed at this stage.

ChIP-seq analyses confirmed the specific binding of ParB to 16 *parS* sequences (as reported before (Donczew et al., 2016)) and showed ParB binding to one additional *parS* site **(Fig. 5A)**. In the wild type strain, ParB clearly bound to the regions adjacent to *parS* sequences extending bidirectionally up to 2.5-3 kb **(Fig. 5A)**, which is consistent with the phenomenon of the ParB spreading when in close clamp conformation. Notably, in the Δ*smc* strain (TM010), ParB binding to DNA was reduced, with the average number of reads two times lower than observed in the wild type strain **(Fig. 4A-C)**. Notably, in the absence of SMC ParB binding adjacent to *parS* sites was almost completely eliminated and comparable to the control Δ*parB* strain **(Fig. 5A)**. However, thorough analyses of ParB-DNA binding over the whole chromosome showed increased non-specific interactions of ParB in the absence of SMC as compared to wild type strain **(Fig. 5D)**. Thus, this indicates that the absence of SMC diminishes ParB association with *parS* and reduces ParB spreading from *parS* sites, however enhances nonspecific ParB association with DNA is. Lowered ParB spreading corresponds to lower complex stability, while its association with non-specific DNA explains the reduced mobility of ParB-HT in the absence of SMC.

**Figure 5.**
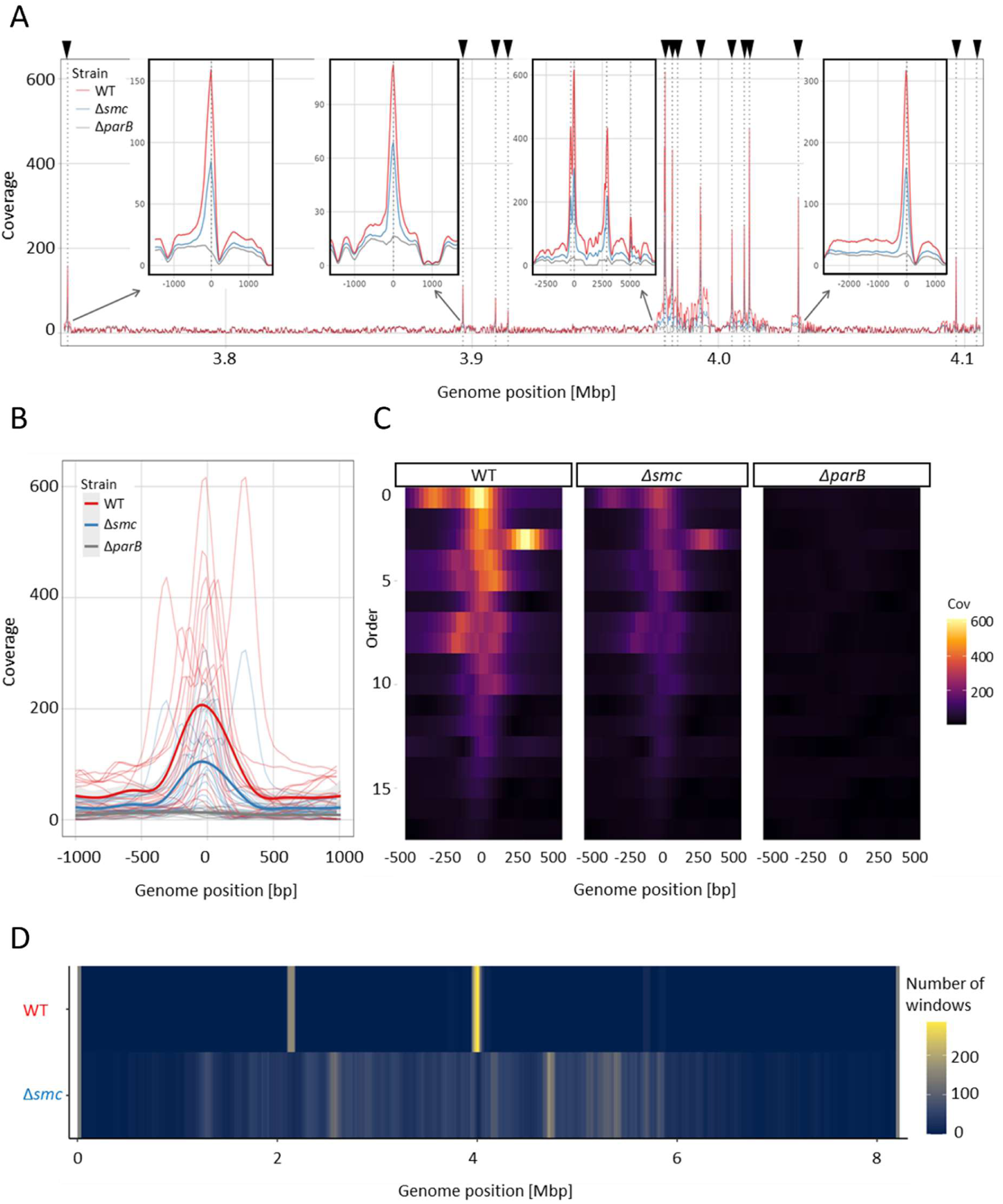
ChIP-seq analyses showing diminished ParB-*parS* binding and spreading in the absence of SMC. ChIP seq was performed using antigen-purified polyclonal ParB antibody and wild type, Δ*parB* (MD020) and Δ*smc* strain (TM010). **A.** ChIP-seq detected binding plotted against the chromosomal region of *S. venezuelae* chromosome containing 17 parS sites in wild type (red), Δ*smc* (blue) and Δ*parB* (grey) strain. Insets show selected *parS* sites. **B**. The average ChIP-seq signal at *parS* sites and neighbouring regions in strain wild type (red), Δ*smc* (blue) and Δ*parB* (grey). **C.** Heatmap of 17 parS sites bound by ParB **D.** Average number of 100 bp long regions with enriched ParB binding in strains: wild type and Δ*smc*. Regions were counted using a 10000 bp rolling window.

### SMC reduces ParB CTPase activity

The spreading of the ParB homologues (*B. subitilis*) was shown to be regulated by CTP binding and hydrolysis (Antar et al., 2021). Therefore, we sought to determine if the reduced *parS*-specific binding and spreading of ParB in the absence of SMC, as indicated by our ChIP-seq data, could be explained by the modulation of its CTP hydrolysis activity by SMC. We undertook a CTP hydrolysis assay measuring ParB CTPase activity in the presence of *parS*-containing plasmid DNA or the presence of the same plasmid DNA with mutated *parS* or in the absence of DNA, as well as in the presence or absence of purified FLAG-SMC **(Fig. S11)**. This revealed that *S. venezueale* ParB CTPase activity increased in the presence of *parS* compared with its activity in the absence of DNA or the presence of mutated *parS*, consistent with data obtained for other ParB homologues (Osorio-Valeriano et al., 2019; Soh et al., 2019) **(Fig. 6)**. Remarkably, the addition of FLAG-SMC to ParB in the presence of *parS*-containing DNA significantly reduced CTP hydrolysis - from approximately 40 μmol of phosphate released by 1 μmol of ParB during 30 min in the absence of FLAG-SMC to 23 μmol of phosphate released in presence of FLAG-SMC **(Fig. 6)**. Thus, SMC decreased ParB CTPase activity, which, is consistent with the observed enhanced ParB spreading along DNA.

**Figure 6.**
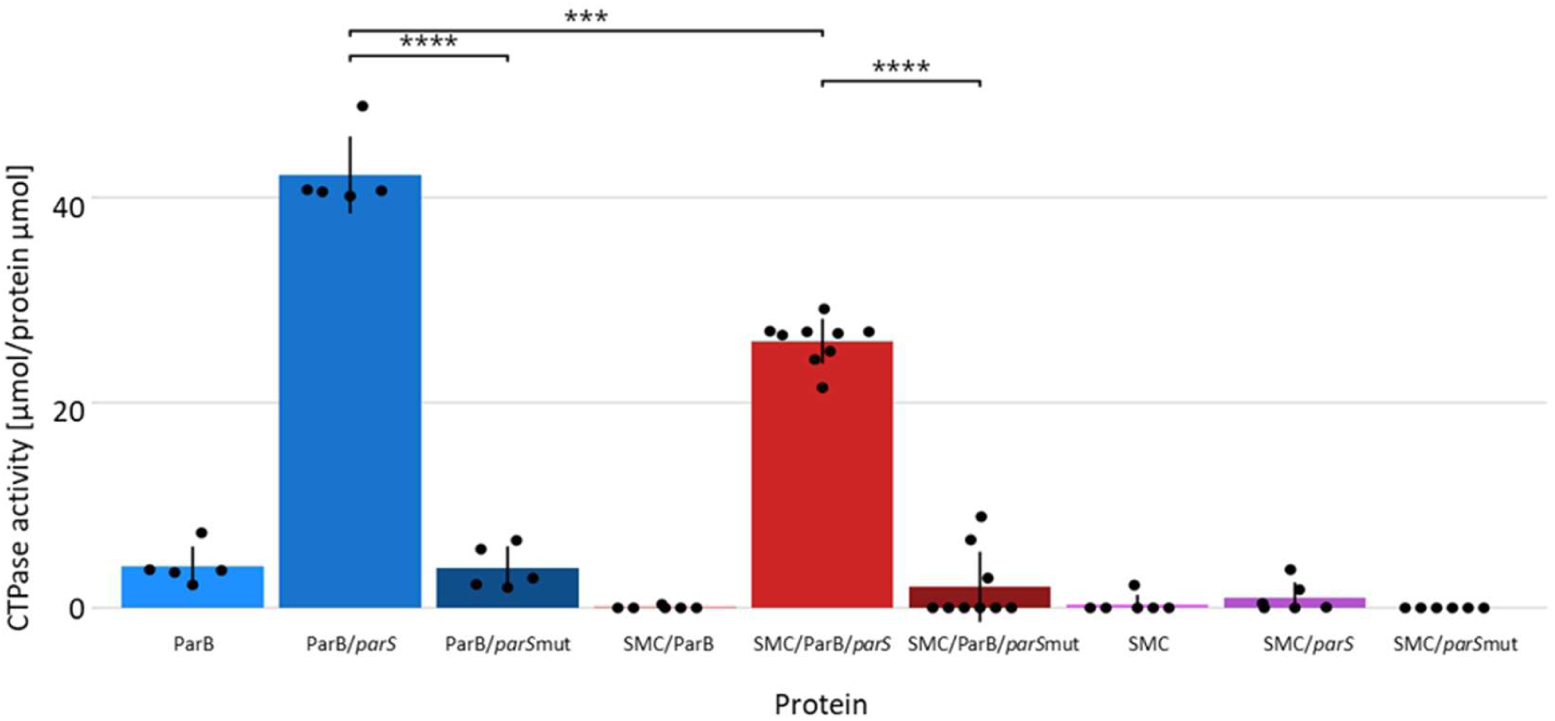
SMC complex reduces ParB CTPase activity. Bar plot showing the results of CTP hydrolysis assay (phosphate detection) for ParB, ParB in the presence of pBSK plasmid with single *parS*, and ParB in the presence of pBSK plasmid with mutated *parS* (ParB-*parS*mut) as compared to ParB in presence of FLAG-SMC, ParB in presence of pBSK plasmid with single *parS* and SMC-FLAG, and ParB in presence of pBSK plasmid with mutated *parS* (ParB-*parS*mut) in presence of SMC-FLAG. Data were obtained from at least 2 independent experimental repeats, each in 2 technical repeats. Two sided t-student test with holm correction for multiple testing was used in statistical analysis.

## DISCUSSION

In summary, our data provide fundamental insight into understanding segregation complex assembly. Our experiments reveal the impact of SMC on ParB complexes, which has not been characterized before. We found that the absence of SMC shortened the lifetime of ParB complexes on DNA and increased their turnover rate in *S. venezuelae* cells, as determined by timelapse microscopy and FRAP analyses, respectively. These data suggest that SMC stabilizes ParB complexes on DNA. Moreover, in cells lacking SMC, diminished binding of ParB to *parS* sites and reduced spreading was demonstrated by ChIP-seq. The diminished ParB spreading in the absence of SMC explains the lowered complex stability. Surprisingly, single molecule tracking showed that the absence of SMC significantly lowered mobility of ParB molecules, which seemingly contradicts lowered complex stability. However, the immobile ParB molecules were dispersed throughout the cell, most likely due to increased non-specific interaction with DNA, which was confirmed by ChIP-seq. Finally, the presence of SMC reduced ParB CTPase activity *in vitro*, further corroborating with SMC promoting ParB spreading. Therefore, our data reveal positive feedback of SMC on the ParB complex stability **(Fig. 7)**.

**Figure 7.**
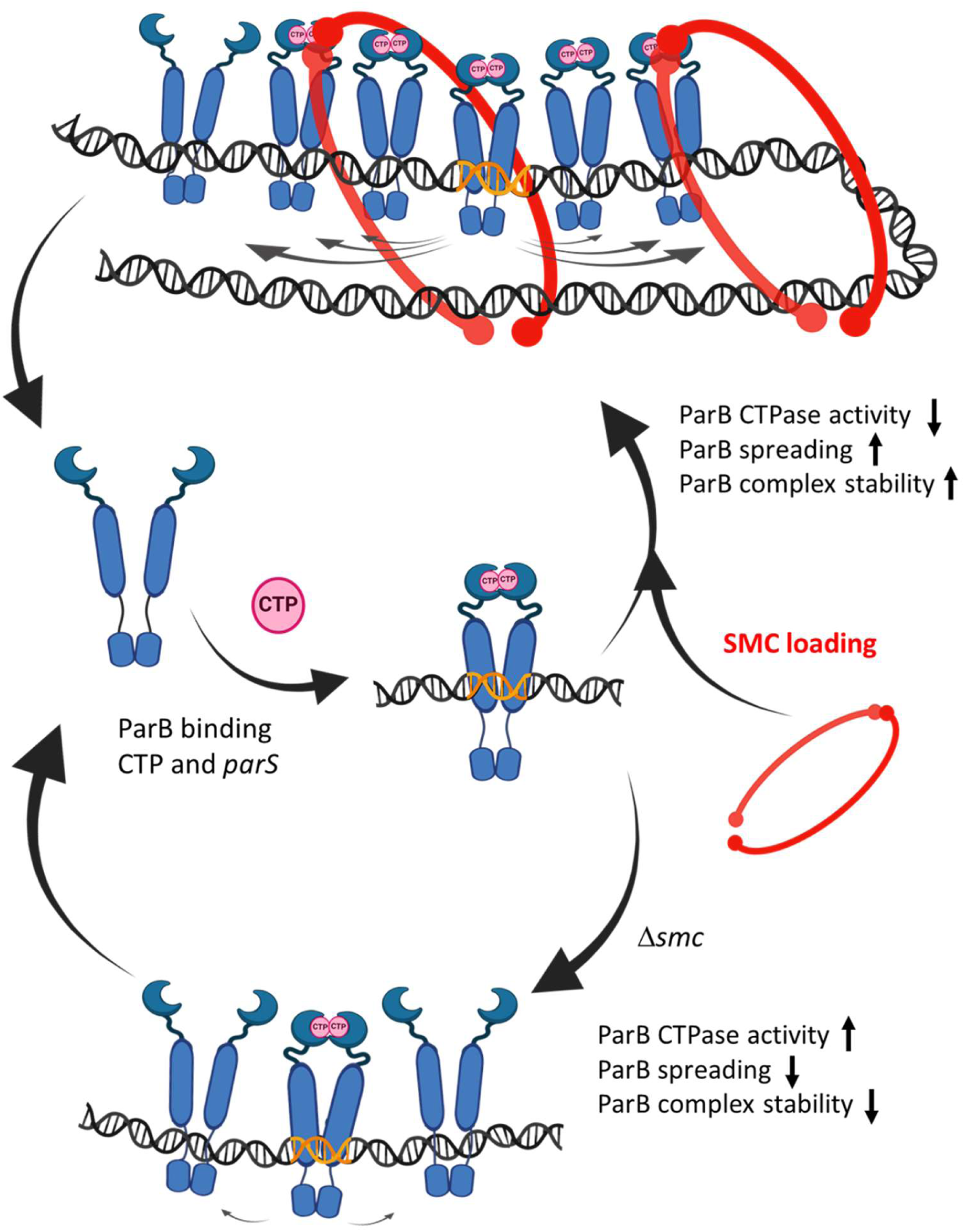
Positive feedback of SMC loading on ParB complex assembly. ParB binds *parS* as the CTP-bound dimer which is followed by a change in the conformation to the closed clamp that slides away from *parS*. SMC loaded on the ParB complex stabilizes ParB in a clamp conformation, preventing clamp opening, inorganic phosphate release and ParB dissociation from DNA and promoting spreading from *parS*. In the absence of SMC ParB clamp on DNA is destabilized and spreading is limited.

It was shown that upon nucleation stage i.e. binding of open ParB dimer to *parS* and CTP, the protein undergoes a conformational change to form a closed clamp that facilitates ParB spreading (Connolley et al., 2023; A. S. Jalal et al., 2021; Osorio-Valeriano et al., 2021). While SMC recruitment to the ParB complex is well documented (Gruber & Errington, 2009; Minnen et al., 2011; Sullivan et al., 2009), the ParB conformation that recruits SMC has not been determined. It remains to be elucidated if SMC is recruited to ParB at the nucleation stage or to the ParB in a closed clamp conformation. Given our finding that SMC promotes ParB spreading, if the former would be the case, SMC could promote ParB change to clamp formation. Consequently, in the absence of SMC, ParB would bind to *parS* less efficiently and spreading would be reduced, as we observe. However, by promoting ParB spreading, SMC might be expected to increase the number of molecules bound to DNA, and a higher number of molecules able to hydrolyse CTP could be reflected in the increased release of inorganic phosphate. That contradicts our observation of less inorganic phosphate released by ParB in the presence of SMC. This leads us to the hypothesis that SMC recruitment rather stabilizes the ParB clamp conformation. The SMC inhibitory effect on CTP hydrolysis by ParB may result from either inhibition of hydrolytic activity per se or inhibition of clamp opening, which would prevent ParB dissociation from DNA and release of hydrolysed CDP and inorganic phosphate. The fact that ParB residues engaged in the interaction with SMC are in close proximity to those responsible for CTP binding and hydrolysis (Bock et al., 2022), supports the conclusion that SMC stabilizes clamp conformation of ParB. SMC may also act at both stages of ParB cycle – condensin could promote ParB clamp closing and stabilize this conformation. Additionally, SMC by extruding DNA loops could enhance binding of ParB molecules to distant regions of DNA, possibly to the other *parS* sites. Such an *in-trans* recruitment of ParB was described earlier (Tišma et al., 2022). Based on available data we cannot determine ParB conformation interacting with SMC.

The surprisingly lowered mobility of ParB molecules in the absence of SMC determined by single-molecule tracking seemingly contradicts the lowered complex stability. Importantly, we observed that ParB mobility was highly elevated in spores when ParB complexes were disassembled, reflecting the ParB release from DNA. This is also consistent with earlier studies showing increased ParB mobility in the absence of *parS* (Osorio-Valeriano et al., 2021). One possible explanation of reduced ParB mobility in the absence of SMC despite its diminished *parS* binding may be ParB ability to form condensates in a non-DNA bound state. The increased dissociation of ParB from *parS* could promote condensate assembly. While some studies suggest that ParB condensates require *parS* sequences, the formation of ParB clusters in the absence of *parS* was also postulated (Babl et al., 2022; Böhm et al., 2020; Guilhas et al., 2020; Zhao et al., 2024). Additionally, the ParB molecules released from *parS* in the absence of SMC may become engaged with other structures such as nucleoid-bound ParA. Our global analyses of ParB binding showed protein association with non-specific DNA, supporting this explanation. Moreover, ParA was reported to influence ParB-SMC interaction, which indicates the crosstalk between both ParB partners (Roberts et al., 2024; Taylor et al., 2021). It should be considered that ParA was shown to be present at the tips of young vegetative cells and along the sporogenic cells (Donczew et al., 2016; Kois-Ostrowska et al., 2016), where we observed lowered ParB mobility in the absence of SMC, but is absent in spores, where SMC had no impact on ParB mobility. Although this hypothesis requires additional investigation, our observation nevertheless provides important insight into ParB regulation by SMC and the crosstalk between SMC and ParA.

In *Streptomyces*, ParB complexes play different roles depending on the growth stage – in sporogenic cells they assist chromosome segregation, while in vegetative cells they anchor chromosomes at the tips of hyphal cells, facilitating efficient tip growth and branching. Our results indicate that SMC stabilizes ParB complexes in sporogenic cells undergoing synchronized cell division. At this stage, ParB complexes have been shown to recruit SMC, which leads to the rearrangement of the chromosome and juxtaposition of the chromosomal arms (Szafran et al., 2021). Intriguingly, we also observed that SMC stabilizes ParB complexes at the early vegetative stage. Little is known about the role of SMC at the earlier stages of growth. The earlier studies showed that in vegetative multigenomic *Streptomyces* cells the chromosomes have predominantly open conformation with limited interarm contacts (Szafran et al., 2021). However, SMC expression is constant through the life cycle of *Streptomyces*, which suggests its involvement in chromosome organization at all stages of growth. It should be noted that ParB complex stability (determined by time-lapse microscopy and FRAP) was much more varied in the presence of SMC compared to its absence. We speculate that the variation of stability may be dependent on the progress or efficiency of SMC loading.

Interestingly, in *S. venezuelae, smc* deletion or *parB* deletion leads to modest chromosome segregation defects. However, both deletions when combined result in a significant increase of anucleate spores which collaborates with the earlier studies on *Streptomyces* (Dedrick et al., 2009; Kois et al., 2009). Interestingly, in *B. subtilis* hydrolysis deficient ParB in the absence of SMC resulted in severe chromosome segregation defects (Antar et al., 2021). The disturbed chromosome segregation in the absence of SMC may be the consequence of disturbed chromosome condensation. In *Streptomyces*, SMC loading and chromosomal arm juxtaposition were shown to contribute to chromosome compaction, which allows regular distribution of the chromosomes along the sporogenic cell (Szafran et al., 2021). Given the hypothesis that the elimination of SMC is likely to affect the ParB-ParA interaction, the elimination of SMC could potentially disturb chromosome segregation by interfering with the ParB-ParA interaction. Thus, SMC may influence chromosome segregation by chromosome compaction, affecting ParB complex stability and its interaction with ParA, however, elucidation of the crosstalk between these three proteins calls for further studies.

In summary, our data provide fundamental insight into the understanding of segregation complex assembly **(Fig. 7)**. Up to now, ParB was also shown to recruit SMC, however, the impact of SMC on ParB complex has not been reported before. Our data prove the positive feedback of SMC on the ParB complex. The presented data set the stage for further mechanistic studies of ParB complex assembly in the presence of SMC. The structural studies are required to fully understand the conformational changes of ParB upon interaction with SMC and to elucidate the crosstalk between three partners ParA-ParB and SMC.

## MATERIALS and METHODS

### Bacterial strains and plasmids

Bacterial strains used in the study are listed in Table S1, and the constructs are listed in Table S2. Genetic manipulations were performed according to standard protocols and followed specific enzyme or kit manufacturer’s recommendations. *E. coli* culture conditions and manipulation methods followed the commonly employed procedures (Russell & Sambrook, 2001). The *S. venezuelae* strains used in this study are derivatives of the NRRL B-65442 strain described as wild type. Strain construction strategies are described in the Supplementary Information. *S. venezuelae* strains were cultured in maltose yeast extract medium (MYM) agar plates or MYM liquid medium as described before (Bush et al., 2013). Conjugation of plasmids and cosmids from *E. coli* ET12567/pUZ8002 to *S. venezuelae* was performed as described previously ((Bush et al., 2013). Exconjugants were selected using MYM medium. *S. venezuelae* growth rate was analysed as described before (Szafran et al., 2021).

### Chromatin immunoprecipitation combined with next-generation sequencing (ChIP-seq)

For chromatin immunoprecipitation, wild type, Δ*smc* and Δ*parB S. venezuelae* (WT, TM010 – three replicates and MD020, two replicates) were cultured in 50 ml liquid MYM medium supplemented with Trace Element Solution (TES, 0.2x culture volume)(Bush et al., 2013) at 30°C with shaking. Cultures were inoculated with 100 ml of spores suspension with OD_600_=10. After 5 h of growth, the cultures were cross-linked with 1% formaldehyde for 30 min followed by blocking for 5 min with 125 mM glycine. Next, half of the culture volume was washed twice with PBS buffer, and the pellet was resuspended in 750 μl of lysis buffer (10 mM Tris-HCl, pH 8.0, 50 mM NaCl, 14 mg/ml lysozyme, protease inhibitor (Pierce)). After 1 h incubation at 37°C for 14 h, 200 μl of zircon beads (0.1 mm, BioSpec products) were added and the samples were disrupted using a FastPrep-24 Classic Instrument (MP Biomedicals; 2 x 45 s cycles at 6 m/s speed with 5-min breaks, during which the lysates were incubated on ice). Next, cell lysates were mixed with 750 μl IP buffer (50 mM Tris-HCl pH 8.0, 250 mM NaCl, 0.8% Triton X-100, a protease inhibitor (Pierce)), and sonicated to shear DNA into fragments ranging from 300 to 500 bp. The samples were centrifuged, and the supernatant was mixed with 2.25μg of affinity-purified rabbit polyclonal anti-ParB antibody (the antibody was purified using *S. venezuelae* ParB immobilized on CN-Br activated Sepharose). Lysates were incubated with antibodies for 14 hours at 4°C, and next 80 µl of Pierce™ Protein A Magnetic BeadsG (Thermo Scientific 88845) were added to the samples. Immunoprecipitation was performed overnight at 4°C. Next, the magnetic beads were washed twice with IP buffer, once with IP2 buffer (50 mM Tris-HCl pH 8.0, 500 mM NaCl, 0.8% Triton X-100, protease inhibitors (Pierce)), and once with TE buffer (Tris-HCl, pH 7.6, 10 mM EDTA 10 mM). After washing, magnetic beads were resuspended in IP elution buffer (50 mM Tris-HCl pH 7.6, 10 mM EDTA, 1% SDS) and incubated overnight at 65°C. Next, the samples were centrifuged, and proteinase K (Roche) was added to the supernatants to a final concentration of 100 µg/ml followed by incubation for 90 min at 55°C. DNA was extracted with phenol and chloroform and subsequently precipitated overnight with ethanol. The precipitated DNA was dissolved in nuclease-free water (10 μl). The concentration of the DNA was quantified using a Qubit dsDNA HS Assay Kit (Thermo Fisher Scientific).

DNA sequencing was performed by NGS Services (Switzerland) using the Illumina ChIP-Seq TruSeq protocol, which included quality control, library preparation and SE sequencing (1×150 bp) of amplified fragments. The bioinformatics analysis was performed using the R package *normr* and the MACS2 programme (Helmuth et al.; Zhang et al., 2008). The mapping of ChIP-seq data was performed using the *Bowtie2* tool (version 2.5.2) (Langmead & Salzberg, 2012). The successfully mapped reads were subsequently sorted using *samtools* (version 1.19.2) (Li et al., 2009). The total number of mapped reads was above 10^6^ on average. Regions specifically bound by ParB were identified using the MACS2 programme. Only regions with fold > 1.75 were considered for further analysis. The unspecific binding of ParB was analyzed using normr package. Reads were counted in a 100 bp long window and diffR function was used to find regions enriched in either wild type or Δsmc strains. Only regions with FDR < 0.001 were considered to be significant.

### FLAG-SMC purification

For the FLAG-SMC pulldown assay *S. venezuelae* TM017 strain was used (Szafran et al., 2021). 1 ml of spore suspension with OD_600_=0.05 was used to inoculate 900 ml liquid MYM medium supplemented with 10 ml TES solution, cultures were grown at 30°C with shaking (200 rpm) for 16 h. The cells were collected by centrifugation (5000 g, 10 min, 4◦C), washed twice with 50 ml of IP buffer (50 mM Tris–HCl pH 8.0, 250 mM NaCl, Triton 0.8%), supplemented with Pierce™ Protease Inhibitor Tablets (Thermo Fisher Scientific, USA) and resuspended in 200 ml of the same buffer. The cells were disrupted by sonication, and the obtained cell lysates were clarified by centrifugation (9 000 g, 20 min, 4°C). The clarified cell lysate was incubated overnight with 200 μl of magnetic beads coated with anti-FLAG® BioM2 antibody (Sigma Aldrich, USA) with constant tube rotation at 4◦C. Next, the magnetic beads were washed thrice with 1 ml of IP buffer. To elute bound protein, the magnetic beads were incubated in 200 μl IP buffer (50 mM Tris–HCl pH 8.0, 250 mM NaCl, Triton 0.8%) supplemented with 3xFLAG peptide (to final concentration 150 μg/μl) at 4°C overnight. Purified FLAG-SMC was stored at −20◦C with 25% glycerol.

### CTP hydrolysis assay

Measurements of ParB protein CTPase activity were performed in Tris-HCl buffer at a final volume of 40 μl with a constant concentration of ParB protein (2 μM) and CTP (4 mM). The CTP hydrolysis was analysed in the presence of pBSK plasmid (15 μM) that contained a single *parS* sequence or scrambled *parS* and/or in the presence of FLAG-SMC (0,2 μM).

The CTP hydrolysis reaction was carried out in transparent 96-well plates for 30 min at 37°C. The released phosphate was detected using the ATPase/GTPase Activity Assay Kit (Merck) according to manufacturer protocol. The standard curve prepared according to kit manufacturer guidelines was used to calculate the amount of phosphate released [μmol] in the analyzed sample. The obtained absorbance results were converted into the number of μmoles of released phosphate per 1 μmol of protein [μmol/μmol of protein]. Statistical analyses were carried out using the Student’s t-test with correction for multiple testing.

### Fluorescent labelling for microscopy analyses

For the fluorescent microscopy analyses, *S. venezuelae* strains were cultured either on glass coverslips (for DNA and cell wall staining, as described before (Jakimowicz et al., 2007) or in liquid cultures (for Halotag labelling). The liquid cultures were conducted in the 2 ml of MYM liquid supplemented with Hoechst 33342 (16,2 nM) for DNA staining, NADA-green (2,5 nM), for peptidoglycan staining) or ligands for the HaloTag protein - TMR *direct ligand* (5 nM) or Janelia Fluor 549 (0,1 nM). All dyes were added to the culture at the time of inoculation. Before the preparation of microscopy samples (i.e. after 5, 17, and 22 hours of growth) the cultures were washed three to eight times with PBS buffer and resuspended in PBS (50-200 ml). Next, the cells were spread on agarose pads prepared using gene frames (Thermo Fisher Scientific, USA) on 1,1 % low gelling temperature agarose (Sigma-Aldrich, USA) in MYM or PBS, for the snapshot and FRAP analyses or single particles tracking, respectively. Samples were covered with an 18 × 18 mm coverslip (High Precision Microscope Cover Glasses, Carl Roth, Germany).

### Time-lapse microscopy

Time-lapse fluorescence microscopy was performed using a previously described protocol (Schlimpert et al., 2016) using the CellAsic Onix microfluidic system using B04A plates (ONIX). The spores were loaded into a microfluidic observation chamber at a pressure of 6 PSI, for 2-15 seconds depending on the density of the spores. After loading the spores, the experiment was conducted at a constant pressure of 3 PSI and a temperature of 30°C. For ParB-HT visualization in the time-lapse microscopy, strains expressing *parB-HT* were cultured in the presence of the Halotag ligand – TMR direct ligand (5 nM) and optionally in the presence of NADA-green (2,5 nM).

The observation was performed using a DeltaVision Elite Ultra High-Resolution microscope equipped with a CoolSnap camera and Olympus PLANApo 100x/1.40 OIL PH3 lens, and the following filter sets: EYFP -FITC (ex., 513/17 nm; em., 548/22 nm), mCherry -mCh (ex 575/25 em 625/45 nm). The images were taken every 10 min for approximately 20 h with an exposure time of 50 ms at 5% transmission for the DIC channel and 80 ms at 50% transmission in the YFP and mCherry channels. The time-lapse movies were analysed using ImageJ Fiji software (Schindelin et al., 2012).

The positions of the fluorescent complexes were identified using custom protocols involving Fiji software and the R (code available at https://github.com/astrzalka/findpeaks). For statistical analysis two-sided t-students test or Wilcoxon test, with an adjustment for multiple testing if necessary, was used.

### Fluorescence recovery after photobleaching

For FRAP experiments the samples on agarose were prepared from 5 h MYM cultures as described above. FRAP analyses were carried out using Stellaris 8 confocal microscope (Leica GmbH, Germany), equipped with HC PL APO CS2 100x/1.40 OIL lens and at a temperature set to 30°C. 540 nm laser line, originating from WLL2 (with a total power > 1.8 mW) set at 80% of the total power, was used to image and bleach the sample. Fluorescent complexes located in proximity to the hyphal tip were selected to bleach. Before bleaching, 3 frames with no interval were acquired with a laser power set to 2.32%. Then the rectangular area covering the fluorescent complex was bleached by acquiring 3 consecutive frames with a laser power set to 95%. In the post-bleach phase, 19 frames were acquired with an interval set to 30 seconds (total post-bleach acquisition time was 9.5 min) and a laser power set to 2.32%. Image analysis was performed in FIJI - using the Stowers set of plug-ins to get the FRAP curves as well as the recovery times and the sizes of mobile fractions (Schindelin et al., 2012) and R packages. In the FRAP experiment, each frame was averaged two times (with the exemption of bleaching phase) and had a size of 168 x 75 pixels. The pixel size was set to 70 nm whereas pixel dwell time was set to 3.2 used.

### Single particle tracking (SPT)

The microscopy samples for SPT were prepared on agarose pads from 5, 17 and 22 h cultures in MYM cultures, as described above. SPT analyses were carried out using the Zeiss Elyra 7 microscope equipped with Alpha Plan-APO 100x/1.46 Oil DIC VIS lens, sCMOS + emCCD and Andor EM-CCD cameras, and following laser lines: 405 (50 mW), 488 (100 mW), 561 (500 mW), 633 nm (500 mW). The imaging was performed with a 561 nm laser with a fixed power range of 80 %. The. From 10,000 to 20,000 frames were collected with an exposure time of 20 ms. The experiment was conducted at a constant temperature of 30°C. Fiji plugin Trackmate (Tinevez et al., 2017), with a maximum linking distance of 0.5 μm, was used for single molecule tracking. Only tracks with at least 4 steps were considered for further analysis. Cell outlines were prepared using the Oufti programme (Paintdakhi et al., 2016). SMTracker was used to analyse ParB mobility in hyphae (Rösch et al., 2018). Square Displacement analysis (SQD) was used to divide analysed ParB molecule tracks into two diffusive populations based on their apparent diffusion constant (D).

## ACKNOWLEDGEMENTS

This work was funded by the Polish National Science Centre: OPUS grant 2018/31/B/NZ1/00614 (to D.J.). We thank Tung Le for the discussion of the results.

## AUTHOR CONTRIBUTION

Conceptualization and experiment design (D.J.), methodology including strains construction and proteins purification (K.P., A.S., J.D-K.), microscopy and data analysis (K.P., A.S. M.M.); ChIP-Seq libraries preparation and data analysis (K.P., A.S.); figures preparation (K.P. and A.S); writing the manuscript (D.J.); manuscript revision and discussion (K.P., A.S., M.S., D.J.).

## DATA AVAILABILITY

The raw ChIP-Seq data, as well as the processed data generated in this study (shown in Fig. 6), have been deposited in the ArrayExpress database (EMBL-EBI) under accession code E-MTAB-14547.

