## Supplementary figures for "SMC modulates ParB engagement in segregation complexes in *Streptomyces*"

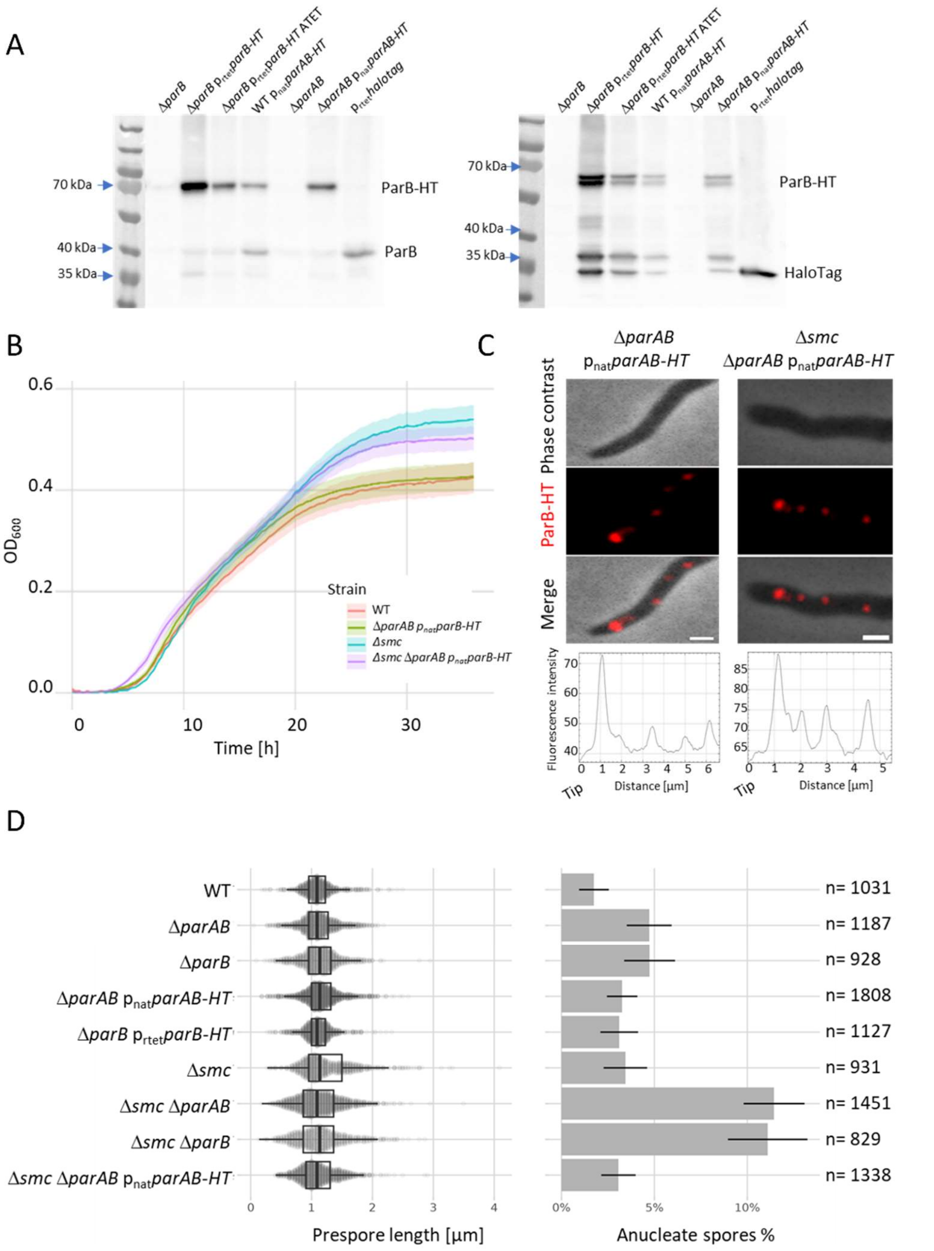


**Figure S1. Verification of ParB-HT producing strains. A.** Western blotting analyses of ParB-HT levels in modified *S. venezuelae* strains (as indicated): Δ*parB* (MD020), Δ*parB* p$_{rtet}$*parB-HT* (KP009) cultured without addition of anhydrotetracycline and Δ*parB* p$_{rtet}$*parB-HT ATET* (KP009) cultured in the presence of 100 ng/ml of anhydrotetracycline, WT p$_{nat}$*parAB-HT* (KP005), Δ*parAB* p$_{nat}$*parAB-HT* (KP006) and WT p$_{rtet}$*halotag* (KP003), Left panel – anti ParB antibody, right panel – anti HaloTag antibody. The positions of ParB, ParB-HT and Halotag are indicated. **B.** Growth curves of wild type *S. venezuelae* strain (WT) and Δ*smc* strain (TM010) compared to growth curve of Δ*parAB* p$_{nat}$*parAB-HT* (KP006) and Δ*smc* Δ*parAB* p$_{nat}$*parAB-HT* (KP007). **C.** Representative images showing ParB-HT complexes stained with TMR in young vegetative cells (5 hours of growth) of Δ*parAB* p$_{nat}$*parAB-HT* (KP006) and Δ*smc* Δ*parAB* p$_{nat}$*parAB-HT* (KP007) strain. TMR -red, phase contrast – gray. Scale bar – 1 μm. The lower panel shows the analyses of fluorescence intensity. **D.** Analysis of septation defects (prespore length) - left panel and chromosome segregation defects (% of anucleate spores) in modified *S. venezuelae* strains. Number of analysed spores (n) is indicated.

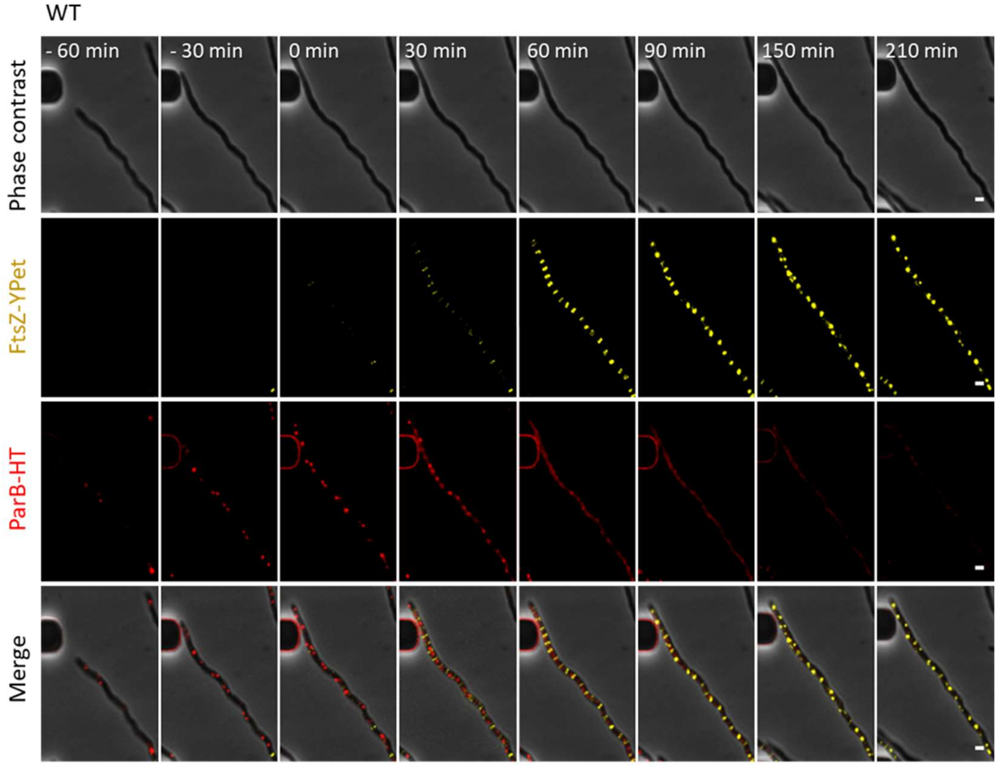


**Figure S2. Time lapse analyses of ParB-HT complexes and septation. A.** Representative images from time lapse analysis of sporogenic development of Δ*parB* $p_{rtet}$*parB-HT ftsZ-ypet* (KP011) strain with fluorescence of ParB-HT stained with Janelia Fluor-549 (red), FtsZ-YPet fluorescence (yellow), phase contrast (grey) and all three channels overlaid. Time 0 is the time of hyphal cell growth arrest, scale bar – 1 μm.

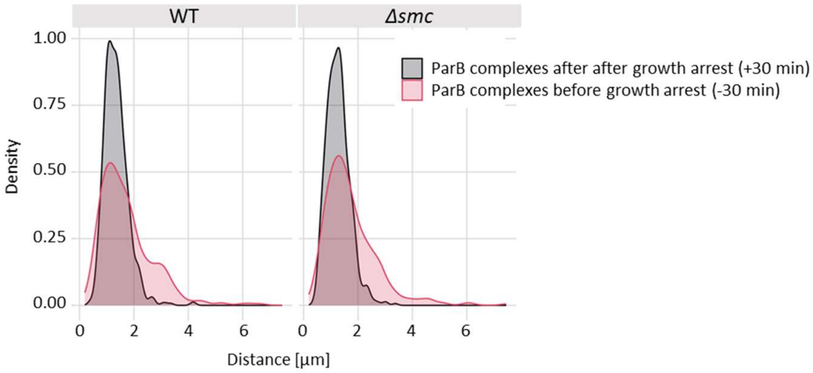


**Figure S3.** Analysis of the distances between the ParB-HT complexes in extending hyphal cells (30 min before growth arrest) and after growth arrest in the Δ*parAB* $p_{nat}$*parAB-HT*, KP006 and in Δ*smc*Δ*parAB* $p_{nat}$*parAB-HT*, KP007 strain. Data were collected in 6 independent experiments for 35 hyphae (before growth arrest) and 73 hyphae (after growth arrest) of KP006 strain and 46 hyphae (before growth arrest) and 78 hyphae (after growth arrest) of KP007 strain.

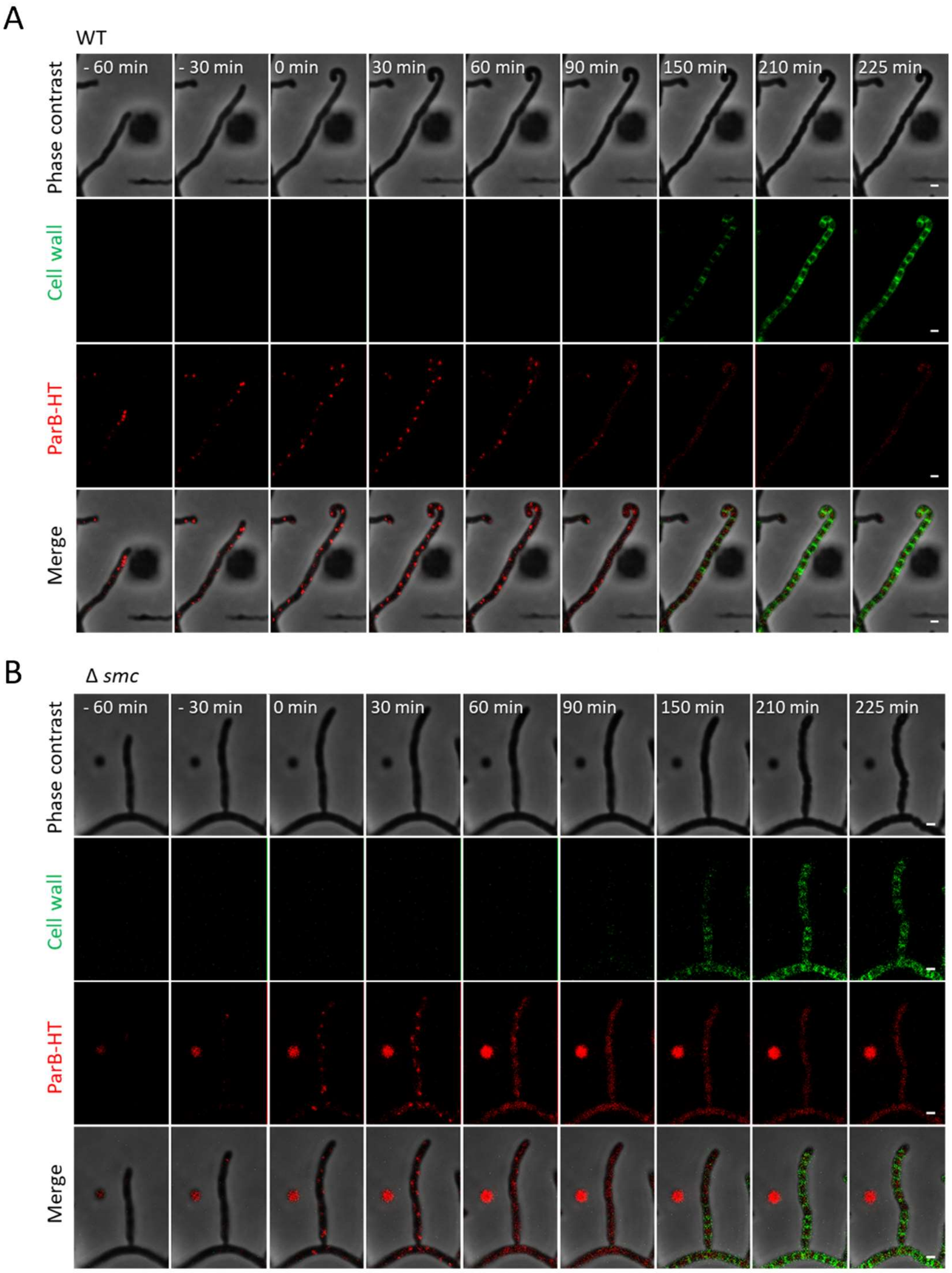


**Figure S4. Time lapse analyses of ParB-HT complexes and septation in wild type and *smc* deletion background A. and B** Representative images from time lapse analysis of sporogenic development of wild type control Δ*parAB* $p_{nat}$*parAB-HT*, KP006 strain **(A)** and of Δ*smc* Δ*parAB* $p_{nat}$*parAB-HT*, KP007 strain **(B)** with fluorescence of ParB-HT stained with Janelia Fluor-549 (red), fluorescence of NADA green-stained septa (green), phase contrast (grey) and all three channels overlaid. Scale bar – 1 μm.

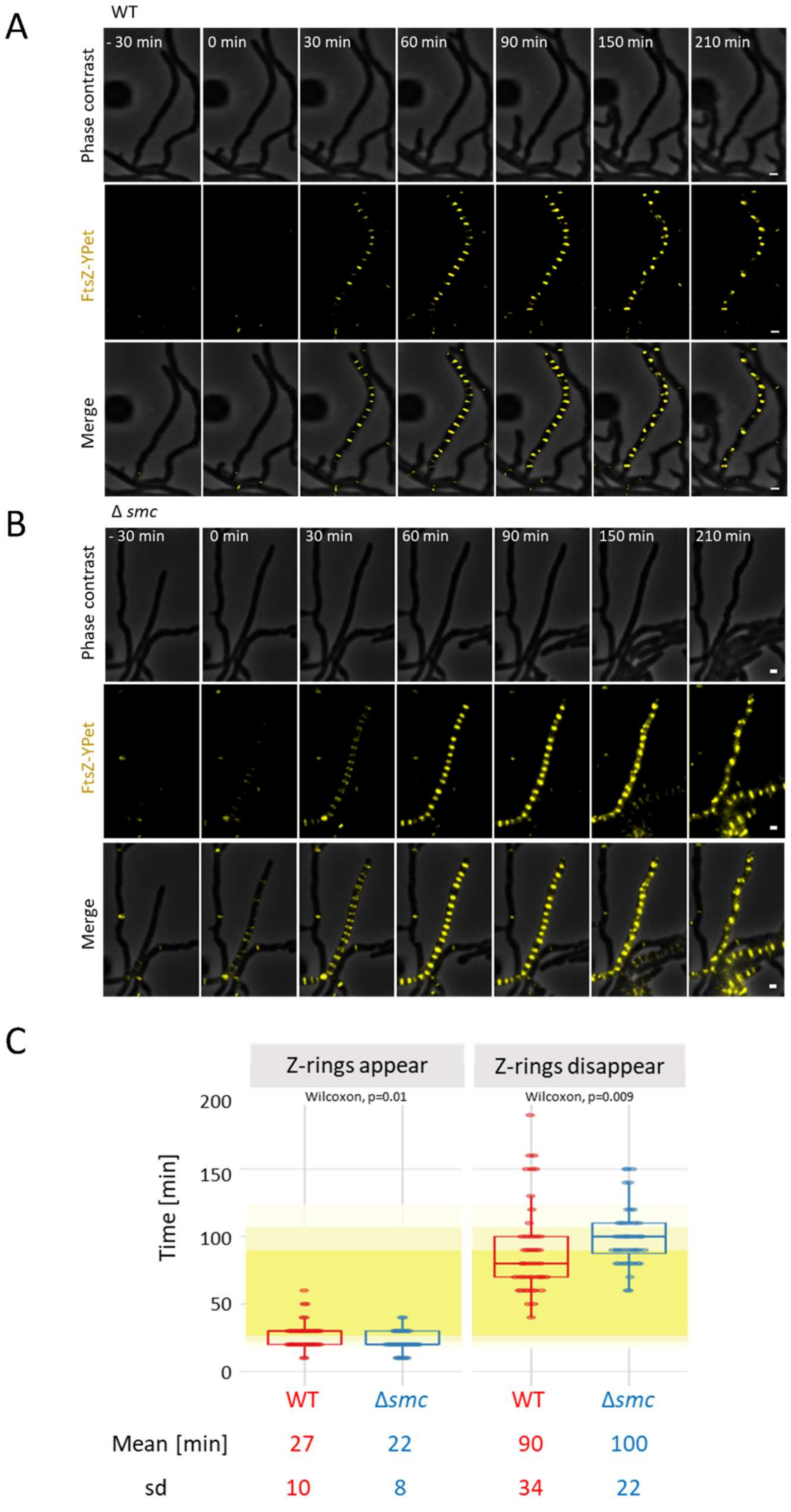


**Figure S5. Time lapse analyses of Z-ring timing in wild type and *smc* deletion background. A and B** Representative images from time lapse analysis of sporogenic development of wt *ftsZ-ypet* (MD100) **(A)** and Δ*smc ftsZ-ypet* strain (TM004) **(B)** showing FtsZ-YPet fluorescence (yellow), phase contrast (grey), and the overlay of two channels. Scale bar – 1 μm. **C.** Analysis of the timing of Z-rings assembly and disassembly in relation to sporogenic cell growth arrest. Yellow shading shows the mean lifetime of Z-rings. Data shown in panel C were collected in 4 independent experiments for 44 hyphae of MD100 and TM004 strains, statistical analyses were performed using two-sided Wilcoxon test.

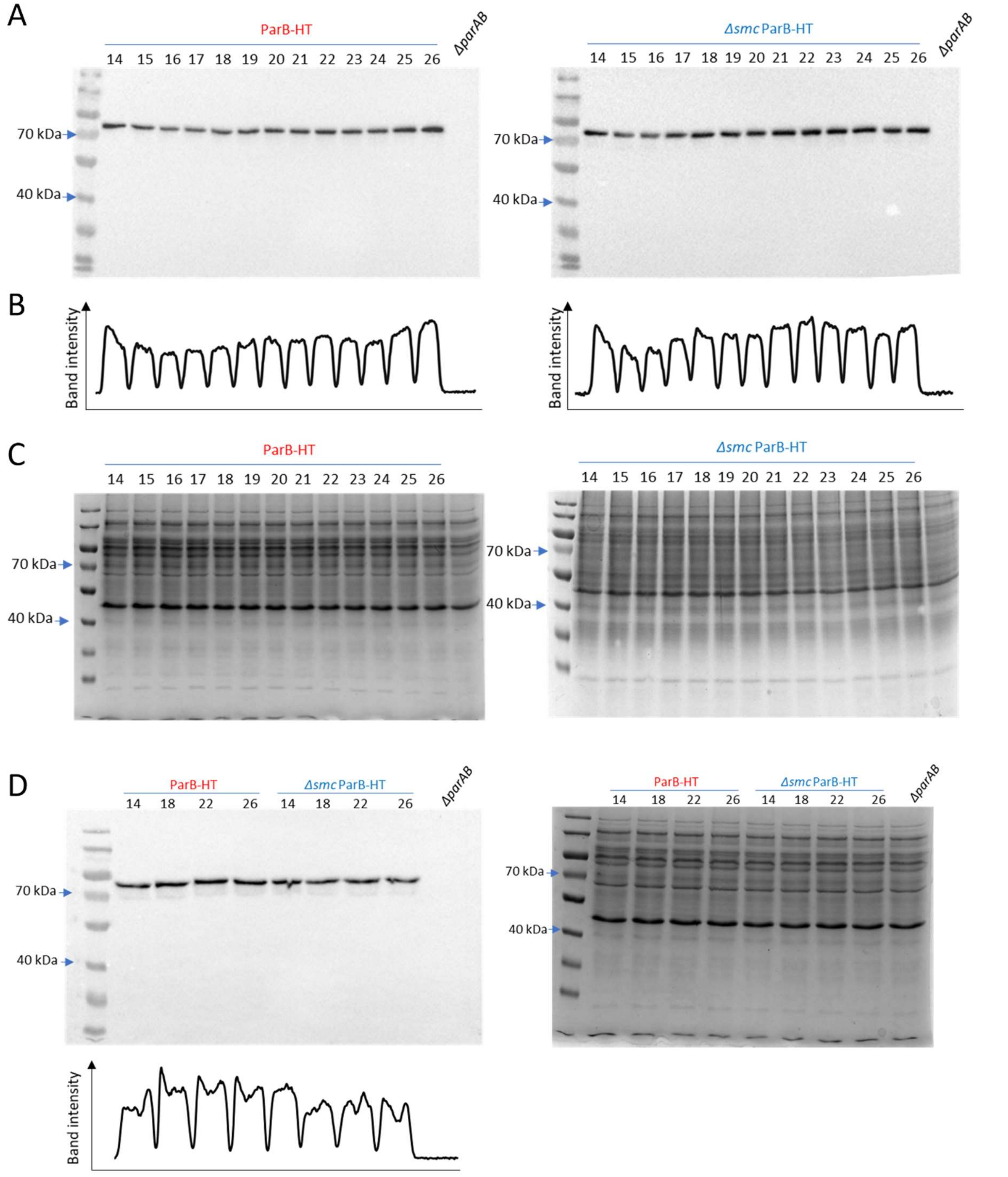


**Figure S6. Levels of ParB-HT in wild type and *smc* deletion background. A** Western blotting detection of ParB-HT in cell lysates of the Δ*parAB* p$_{nat}$*parAB-HT*, KP006 and Δ*smc*Δ*parAB* p$_{nat}$*parAB-HT,* KP007 strain (as indicated) from different time points during sporogenic development (14 h to 26 h of culture). Anti HaloTag antibody. **B**. Analysis of band signal intensity **C**. Loading control – cell lysates analysed in Western blotting shown in panel A in SDS PAGE gel stained with Coomasie brilliant blue. D. Comparison of ParB-HT levels in Δ*parAB* p$_{nat}$*parAB-HT*, KP006 and Δ*smc*Δ*parAB* p$_{nat}$*parAB-HT,* KP007, right panel – loading control, bottom panel analysis of band signal intensity.

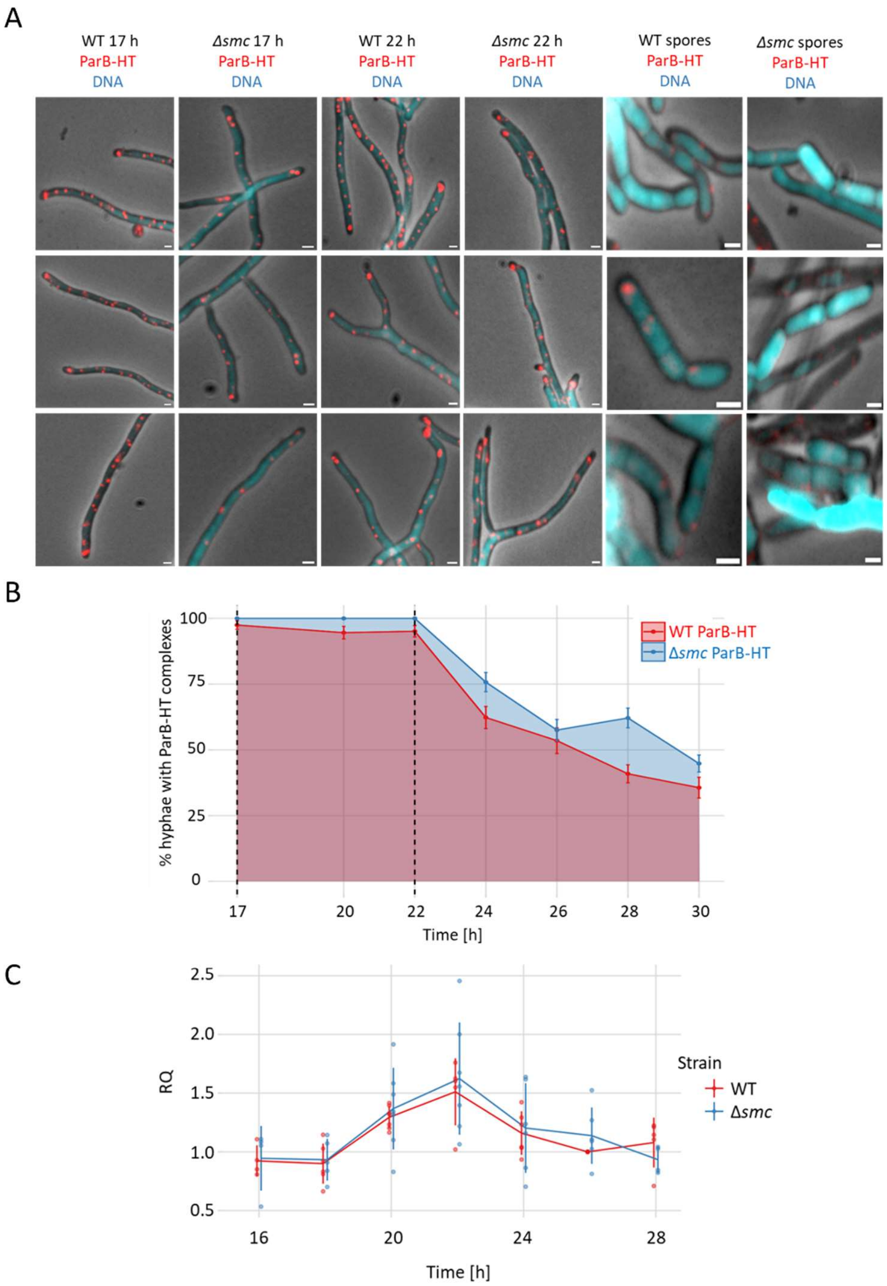


**Figure S7. The progress of sporogenic development of strains producing ParB-HT in wild type and Δ*smc* background determined by the percentage of hyphae showing ParB complexes. A.** Representative images of hyphal cells of the wild type control and Δ*smc* strains (Δ*parAB* p$_{nat}$*parAB-HT*, KP006 and Δ*smc*Δ*parAB* p$_{nat}$*parAB-HT* KP007, respectively) from three time points of sporogenic development. ParB-HT was stained with TMR Direct Ligand (red) and DNA was stained with Hoechst 33342 (blue) overlaid on phase contrast (grey). Scale bar – 1 μm. **B.** The percentage of hyphae showing ParB complexes (either regularly or irregularly spaced) in wild type control and Δ*smc* strains (Δ*parAB* p$_{nat}$*parAB-HT*, KP006 (350-802 hyphae) and Δ*smc*Δ*parAB* p$_{nat}$*parAB-HT* KP007 (187-915 hyphae), respectively) at subsequent time points of sporogenic development. Error bars show 95% confidence intervals. **C**. Efficiency of chromosome replication determined by marker frequency analysis – *ori:ter*

ratio during sporogenic development of wild type control and $\Delta$*smc* strains ($\Delta$*parAB* $p_{nat}$*parAB-HT*, KP006 and $\Delta$*smc*$\Delta$*parAB* $p_{nat}$*parAB-HT* KP007, respectively). Error bars show standard deviation.

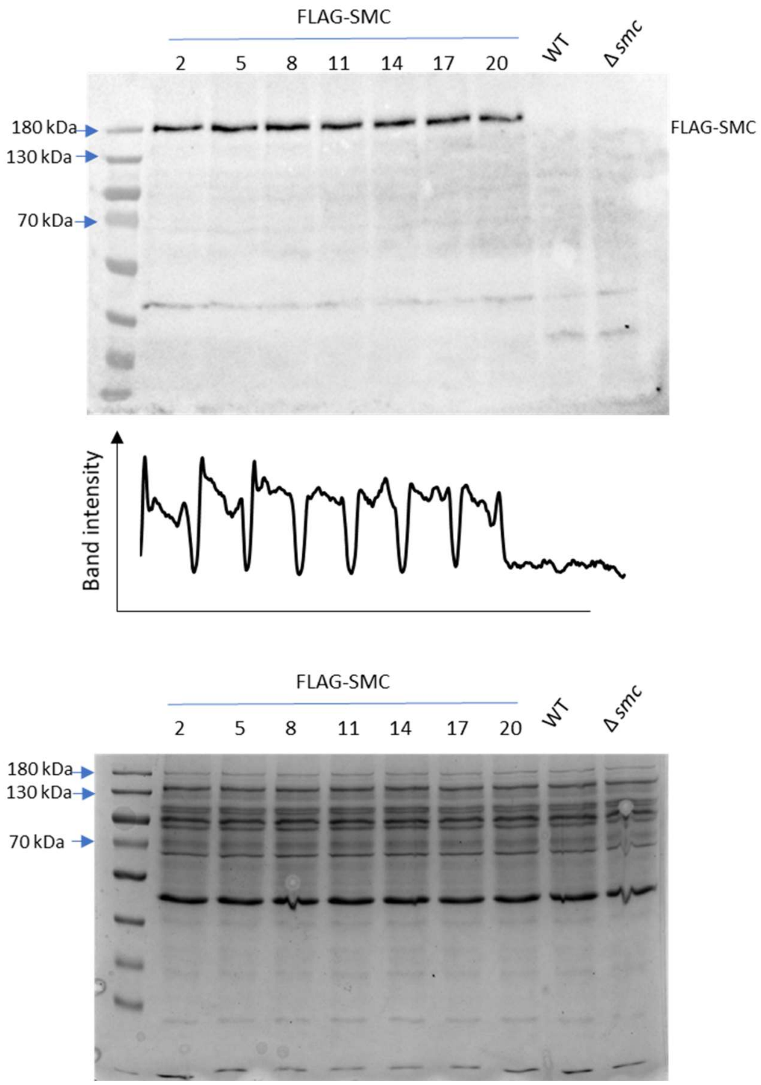


**Figure S8. FLAG-SMC levels during life cycle of *S. venezuelae*.** Western blotting detection of FLAG-SMC in cell lysates of the TM017 strain from different time points of the life cycle (2 h to 20 h of culture, as indicated) using anti FLAGtag antibody. Middle panel shows analysis of band signal intensity, bottom panel shows loading control. The wild type strain (WT) and $\Delta$*smc* strain (TM10) serves as the negative controls.

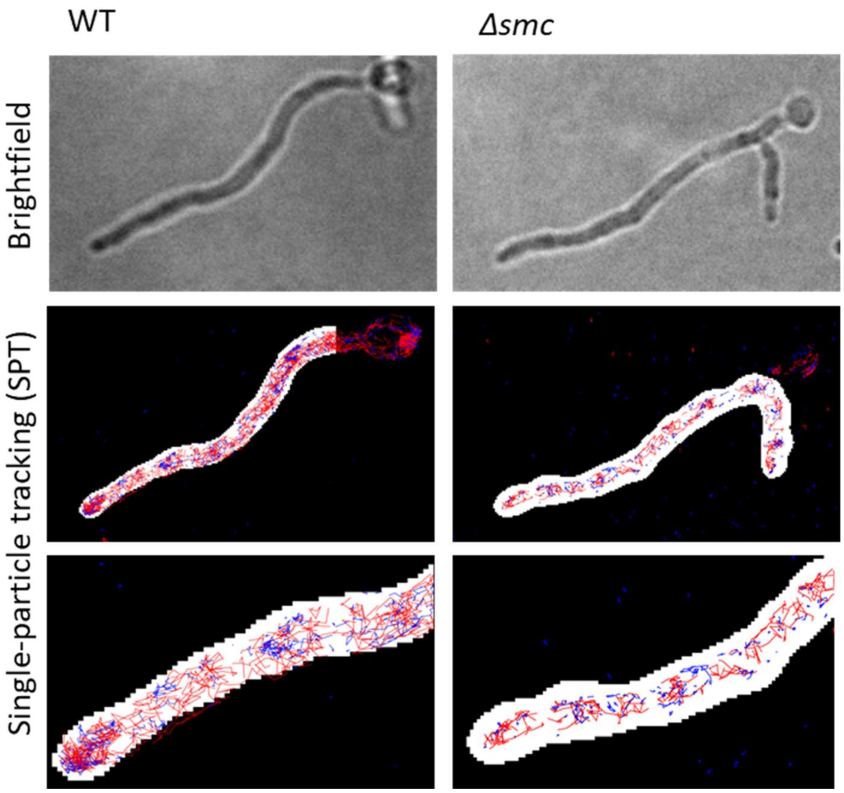


**Figure S9. Single molecule tracking of ParB-HT.** Representative hyphae of the wild type control and Δ*smc* strains (Δ*parAB* p$_{nat}$*parAB-HT*, KP006 and Δ*smc*Δ*parAB* p$_{nat}$*parAB-HT* KP007, respectively) Top images - brightfield images of analysed hyphae, middle and bottom images - ParB-HT tracking labelled accordingly to mean speed – red: fast moving, blue: slow moving molecules. Bottom images are scaled up.

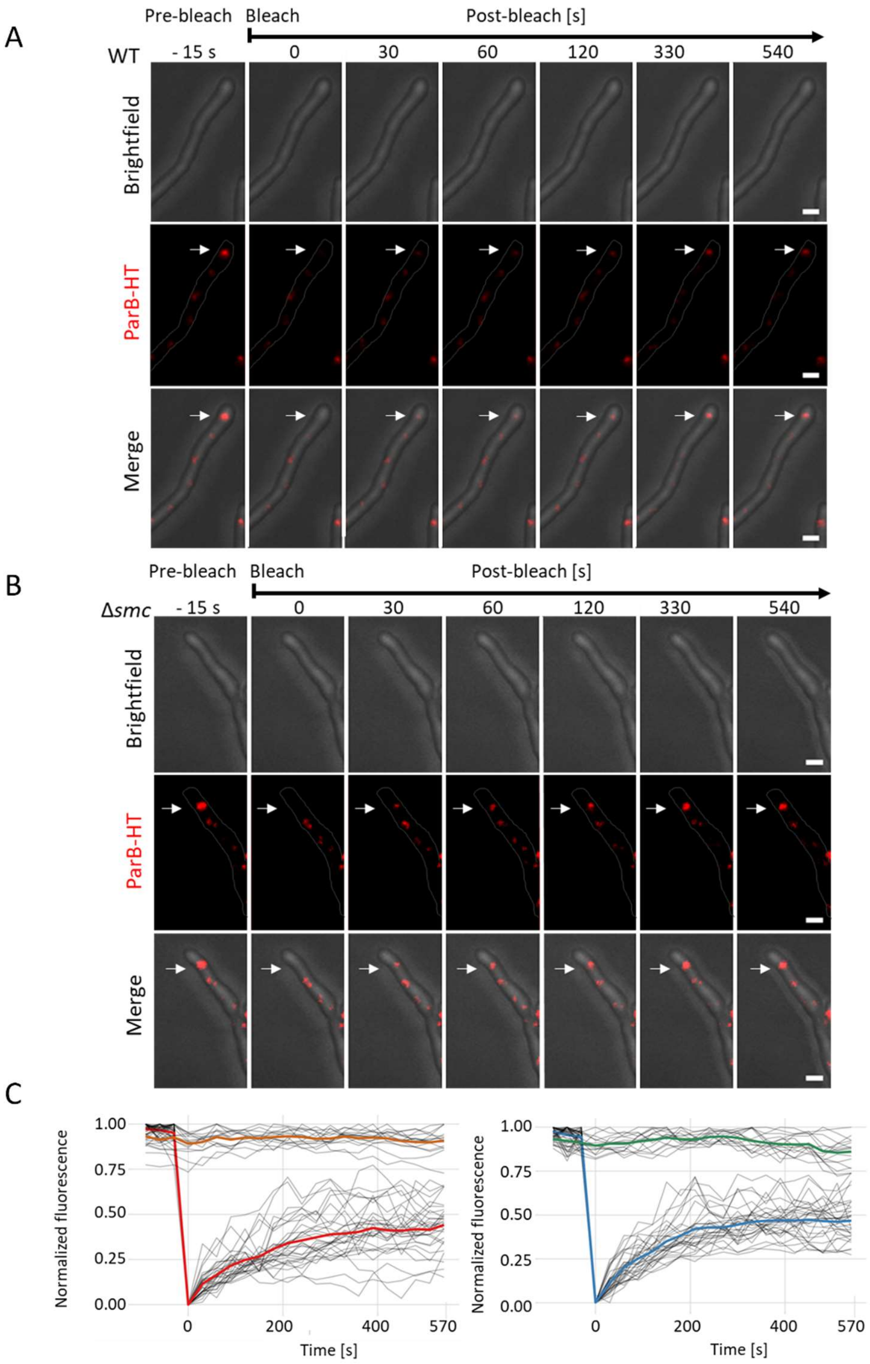


**Figure S10. FRAP analysis ParB-HT complexes in early vegetative cell of the wild type and Δ*smc* background. A** Representative images showing the photobleaching of the ParB-HT complex stained with Janelia Fluor-549 in young vegetative cells of the wild type control and Δ*smc* strains (Δ*parAB* $p_{nat}$*parAB-HT*, KP006 and Δ*smc*Δ*parAB* $p_{nat}$*parAB-HT* KP007, respectively), ParB-HT fluorescence (red) brightfield channel (grey) and overlay of both channels **B.** Fluorescence recovery analysis. Intensity of normalized bleached fluorescence plotted against the time of analyses compared to control fluorescence signal (not bleached) in the wild type control (left panel) and Δ*smc* strain (right panel). 22 complexes were analysed in the wild type and Δ*smc* background (Δ*parAB* $p_{nat}$*parAB-HT*, KP006 and Δ*smc*Δ*parAB* $p_{nat}$*parAB-HT,* KP007).

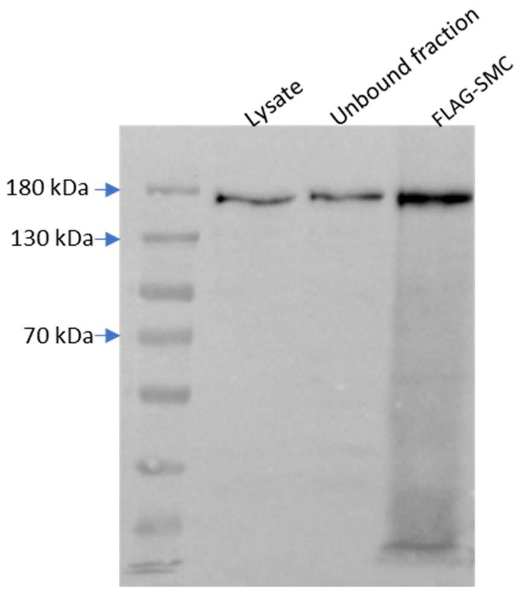


**Figure S11. Western blotting analysis of FLAG-SMC purification from *S. venezuelae*.** FLAG-SMC was immunoprecipitated from 20-hour culture of TM017 strain using magnetic beads coated with anti-FLAG® BioM2 antibody and released with 3XFLAG peptide. FLAG-SMC was detected using anti-FLAG antibody in the TM017 cell lysate, fraction not bound to magnetic beads and fraction released from magnetic beads with 3XFLAG peptide (FLAG-SMC).
