## Supplementary information for "SMC modulates ParB engagement in segregation complexes in *Streptomyces*"

### Table S1. Bacterial strains used in this study

| Name | Description | Source |
|---|---|---|
| ***E. coli*** | | |
| DH5α | F-, Φ80dlacZM15, recA1, *endA1, gyrA96, thi-1,* *hsdR17, (rk-, mk+), supE44, relA1, deoR, (lacZY*AargF) *U169* | Lab stock |
| ET12567/pUZ8002 | Szczep *E. coli* ET12567*: dam13::TN9, dcm6, hsdM, hsdR, recF134, zjj201::TN10, galK2, galT22, ara14, lacY1, xyl5, leuB6, thi1, tonA31, rpsL136, hisG4, tsx78, mtli, glnV44*, F-, ($Cml^R$ ) Plazmid pUZ8002: tra, $Kan^R$,RP4 23 | (Kieser et al., 2000) |
| ***S. venezuelae*** | | |
| WT | Wild-type strain *Streptomyces venezuelae* w bazie *NRRL culture collection* deposited under NRRL number B-65442, by NZ_CP018074.1 | Kind gift from Prof. Mark Buttner John Innes Centre, Norwich, UK (Gomez-Escribano et al., 2021) |
| TM010 (Δ*smc*) | NRRL B-65442 *Δsmc* :: scar | (Szafran et al., 2020) |
| TM017 (*FLAG-smc*) | *smc* :: *FLAG-smc* | (Szafran et al., 2020) |
| TM004 (Δ*smc*; *ftsZ-ypet*) | TM010 attBΦC31 :: pKF351 *ftsZ-ypet* , $Apr^R$ | (Szafran et al., 2020) |
| MD002 (*ΔparB::apra*) | NRRL B-65442 *Δpar ::apra*, $Apr^R$ | (Donczew et al., 2016) |
| MD020 (Δ*parB*) | NRRL B-65442 *ΔparB*::scar | (Donczew et al., 2016) |
| MD030 (Δ*parAB*) | NRRL B-65442 *ΔparAB*::scar | (Donczew et al., 2016) |
| MD100 (*ftsZ-ypet*) | NRRL B-65442 attBΦC31::pKF351, $Apr^R$ | (Donczew et al., 2016) |
| KP003 (WT $p_{rtet}$*halotag*) | NRRL B-65442 attBϕBT1::pKP03 (pSS170 $p_{rtet}$*halotag*), $Hyg^R$ | This work |
| KP004 (Δ*smc*Δ*parAB*) | TM010 Δ*parAB*::*apra*, $Apr^R$ | This work |
| KP020 (Δ*smc*Δ*parB*) | TM010 *ΔparB::apra,* $Apr^R$ | This work |
| KP005 (WT$p_{nat}$*parAB*-HT) | NRRL B-65442 attBϕBT1::pKP05 (pSS170 $p_{nat}$*parAB*-HT), $Hyg^R$ | This work |
| KP006 (Δ*parAB*$p_{nat}$*parAB*-HT) | MD030 attBϕBT1::pKP05 (pSS170 $p_{nat}$*parAB*-HT), $Hyg^R$ | This work |

| | | |
|---|---|---|
| KP007 (Δ*smc*Δ*parAB*$p_{nat}$*parAB*-HT) | KP004 attBϕBT1::pKP05 (pSS170 $p_{nat}$*parAB*-HT), $Apr^R$, $Hyg^R$ | This work |
| KP008 (WT $p_{rtet}$*parB*-HT) | NRRL B-65442 attBϕBT1::pKP08 (pSS170 $p_{rtet}$*parB*-HT), $Hyg^R$ | This work |
| KP009 (Δ*parB* $p_{rtet}$*parB*-HT) | MD020 attBϕBT1::pKP08 (pSS170 $p_{rtet}$*parB*-HT), $Hyg^R$ | This work |
| KP010 (Δ*parB::apra* $p_{rtet}$*parB*-HT) | MD002 attBϕBT1::pKP08 (pSS170 $p_{rtet}$*parB*-HT), $Apr^R$, $Hyg^R$ | This work |
| KP011 (Δ*parB* $p_{rtet}$*parB*-HT *ftsZ-ypet*) | KP009 attBϕC31::pKF351, attBϕBT1::pKP08 (pSS170 $p_{rtet}$*parB*-HT), $Apr^R$, $Hyg^R$ | This work |

**Table S2. Constructs used in this study**

| **Name** | **Description** | **Source** |
|---|---|---|
| pSS170 (pIJ10770) | pMS825 derivative, integrative vector attBΦBT1, *ori* pBR322, *oriT* (RP4) $Hyg^R$ | Kind gift from dr S. Schlimpert, John Innes Centre, Norwich, UK (Bush et al., 2017) |
| pSS170 *halotag* | pSS170 derivative, carrying *halotag* gene, attBΦBT1, *ori* pBR322, *oriT* (RP4) $Hyg^R$, | (Duława-Kobeluszczyk et al., 2024) |
| pSS170$p_{erm}$ *halotag* | pSS170 derivative, carrying *halotag* gene under the control of the $p_{erm}$ promotor, attBΦBT1, *ori* pBR322, *oriT* (RP4), $Hyg^R$, | (Duława-Kobeluszczyk et al., 2024) |
| pTC-28S15-0X $p_{smyc}$*tetR*$_{rv}$$p_{tcp}$*topA*$_{Ms}$ | A non-integrative plasmid carrying *M. smegmatis topA* gene under the control of the $p_{tcp830}$ promoter and the *tetR*$_{rv}$ gene under the control of the $p_{myc}$ promoter (Klotzsche et al., 2009), *ori* pBR322, $Kan^R$, | (Szafran et al., 2018) |
| pKP03 (pSS170 $p_{rtet}$*halotag*) | pSS170 derivative, carrying *halotag* under the control of the $p_{tcp830}$ promoter with an $RBS_{topA,}$ and the *tetR*$_{rv}$ gene under the control of the $p_{s14}$ promoter (Klotzsche et al., 2009), attBΦBT1, *ori* pBR322, *oriT* (RP4), $Hyg^R$ | This work |
| pKF351 | pIJ6902 derivative, carrying *ftsZ-ypet* gene under the control of a native promoter $p_{ftsZ}$, attBΦC31, *oriT* (RP4), $Apr^R$ | (Donczew et al., 2016) |
| pKP05 (pSS170 $p_{nat}$*parAB-HT*) | pSS170 derivative, carrying *S. venezuelae parAB* genes under the control of the native *parAB* promoter, with *parB* in fusion with the *halotag* gene, attBΦBT1, *ori* pBR322, *oriT* (RP4), $Hyg^R$ | This work |
| pKP08 (pSS170 $p_{rtet}$*parB-HT*) | KP03 derivative, carrying *parB-halotag* gene under the control of the $p_{tcp830}$ | This work |

| | | |
|---|---|---|
| | promoter, and an $RBS_{topA,}$ and the *$tetR_{rv}$* gene under the control of the $pS_{14}$ promoter attBΦBT1, *ori* pBR322, *oriT* (RP4), $Hyg^R$ | |
| pCRISPR $p_{tcp}$ $RBS_{topA}$ | Derivative of pCRISPR (Tong et al., 2015), carrying *dcas9* gene controlled by $p_{tcp}$ promoter and $RBS_{topA}$, ApmR, *ori* pBR322, *oriT* (RP4) | Gongerowska-Jac M. , unpublished |
| Sv-4-A09 Δ*parB::apra* | SuperCos-1 cosmid Sv-4-A09 derivative, *parB*::*apr-oriT-FRT*, $Amp^R$, $Kan^R$, $Apr^R$ | (Donczew et al., 2016) |
| Sv-4-A09 Δ*parAB::apra* | SuperCos-1 cosmid Sv-4-A09 derivative, *parAB*::*apr-oriT-FRT* cassette, $Amp^R$, $Kan^R$ $Apr^R$ | (Donczew et al., 2016) |
| pBSK*parS* | pBSKc carrying a fragment of *parAB* promoter from *S. coelicolor* (365 bp), containing a single *parS* site | This work |
| pBSK*$parS_{mut}$* | pBSKc carrying a fragment of *parAB* promoter from *S. coelicolor* (365 bp), containing a mutated *parS* site | This work |

**Table S3. Oligonucleotides used in this work**

| Oligonucleotide | Sequence 5' → 3' | Application |
|---|---|---|
| KP_38Fw | GTACCGCGGATCGTGCTCATGTTCTCTCCCTTGAATTCTAAT | Construction of pKP03 (pSS170 $p_{rtet}$*halotag*) |
| KP_38Rv | TCGATGATCATATGAGAGAATCTAAGCTT TACGTAGACCTACGCCTTGACCTTG | |
| KP_39Fw | ATATGATCATCGATTCGCGACTTAAGCCTAGGCCACCTGAC CGCACGCCCGCAA | |
| KP_39Rv | TCGGATCCATCGTTATTCCTAGGCATATGTCGCTCTTCTCTCC GGGAT | |
| KP_37Fw | AAGGGGATGATAAGTTTATCAAGCTTCCATGGTCATTAGGA GCCGCTCT | |
| KP_37Rv | TTAGAATTCAAGGGAGAGAACATGAGCACGATCCGCGGTA C | |
| KP_38.2Rv | TCATATGAGAGAATCTAAGCTTTACGTAGACCTACGCCTTG AC | |
| KP_37BHTRv | GCTCACTCAACTGGATCCCCTCCTGTCGCTCTTCTCTCCGGG ATA | Construction of pKP05 (pSS170 $p_{nat}$*parAB*-HT) and pKP08 (pSS170 $p_{rtet}$*parB*-HT) |
| KP_66Fw | GATTCGCGACTTAAGCATTCTAGAGGGGATCGTCGATCCGC tctcc | |
| KP_43Rv | TCGGATCCATCGTTATTCCTAGGCATCTCGAGGCCCTCGGC GTCCTCGGCGTT | |
| KP_43BHTFw | TATCCCGGAGAGAAGAGCGACATGGAGGGGATCCAGTGA GTGAGCGA | |
| pSSseq_Fw | AGGATCTTCACCTAGATCCTTTTGGT | Amplification of the pSS170 plasmid insert |
| pSSseq_Rv | GCCAGTGGTATTTATGTCAACACCGC | |

| | | |
|---|---|---|
| pSv_parABkont3_Fw | CCCGCGAGATCGCCCTG | Verification of *parA* and/or *parB* deletion |
| pSv_parABkont4_Rv | ATCCAGGCCTCCTTCTCCA | |
| gyr2_Fw | GCTCCGCTATCACAAGATCA | Marker frequency analysis: amplification of a fragment of the S. *venezuelae chromosome* 3,958,240 – 3,958,322 |
| gyr2_Rv | ACAGGAAGGTCAGCAGCAG | |
| arg3_Fw | CACCTGCGGATCTACAAGC | Marker frequency analysis: amplification of a fragment of the S. *venezuelae chromosome* 1,320,274 – 1,320,351 |
| arg3_Rv | CCACTCCGACATCTCCTTG | |
| parA_fw | CGCAAGCTTGCGCCGCGGCCGACCCCCGC | Amplification of the *parAB* promoter region from *S. coelicolor* |
| parAB_rv | CCGGATCCGACCCGGGTCTGCTCGGGTCGC | |
| parS_mut fw | CGGATGTTTCTAGGGAAACACCGC | Mutation of the *parS* site within the *parAB* promoter region from *S. coelicolor* |
| parS_mut rv | GCGGTGTTTCCCTAGAAACATCCG | |

### SUPPLEMENTARY METHODS

#### Construction of the KP004 (Δ*smc*, Δ*parAB*) and KP020 (Δ*smc*, Δ*parB*) strains

To construct a KP004 (∆*smc*∆*parAB*) and KP020 (∆*parB*∆*smc*), TM010 (Δ*smc*) strain (Szafran et al., 2021) was modified. The exchange of *parB* or *parAB* genes for a cassette containing the apramycin resistance gene and the *oriT* sequence was performed by intergeneric conjugation of the TM010 strain with *E. coli* ET12567/pUZ8002 containing the modified cosmid Sv-4-A09 *ΔparB* or Sv-4-A09 Δ*parAB*. The exconjugants sensitive to kanamycin and resistant to apramycin were selected. Obtained clones were verified with PCR (with pSv_parABkont3_Fw/pSv_parABkont4_Rv primers) and PCR products were sequenced. Additionally, strains were verified using Western blotting with anti-ParB.

#### Construction of *S. venezuelae* KP003 (WT $p_{rtet}$*halotag*) strain

To construct the pKP03 (pSS170 $p_{rtet}$*halotag*), a 277 bp fragment encompassing $p_{tcp}$ promoter was amplified using KP_39Fw/KP_39Rv primers and pCRISPR-$p_{tcp}$-$RBS_{topA}$ plasmid as the template. The resulting fragment was cloned by the SLIC reaction to the pSS170*halotag* vector digested with the XmaJI restriction enzyme. The reaction mixture was used to transform DH5α cells and hygromycin resistant transformants were selected. Next, the *tetRrev* gene (722 bp) was amplified using KP_37Fw/KP_37Rv primers and pTC-28S15-0X $p_{smyc}$*tetR*$_{rv}$$p_{tcp}$*topA*$_{Ms}$ as a template. At the same time, the fragment encoding $p_{S14}$ promoter (120 bp) was amplified using KP_38Fw/KP_38Rv primers and pCRISPR-$p_{tcp}$-$RBS_{topA}$ as a template. An overlap PCR using KP_37Fw/KP_38.2Rv primers amplified both fragments: *tetRrev* gene and $p_{S14}$ promoter, delivering an 801 bp product. The PCR product was cloned by SLIC to the pSS170-$p_{tcp}$halotag vector digested with restriction enzyme HindIII. The reaction mixture was used to transform DH5α cells and hygromycin resistant transformants were selected. The obtained pKP03 (pSS170 $p_{rtet}$*halotag*) vector was verified by PCR (using pSSseq_Fw/pSSseq_Rv primers) followed by sequencing of PCR product.

The pKP03 (pSS170 $p_{rtet}$*halotag*) plasmid was introduced into wild type *S. venezuelae* (NRRL B-65442) by conjugation with *E. coli* ET12567/pUZ8002 followed by selection of hygromycin-resistant exconjugates. The obtained KP0003 strain was verified by PCR reaction (pSSseq_Fw/pSSseq_Rv primers) and Western blotting using an anti-HaloTag antibody.

**Construction of *S. venezuelae strains* producing ParB-HT: KP005 (WT-$p_{nat}$*parAB-HT*), KP006 (Δ*parAB*$p_{nat}$*parAB-HT*) and KP007 (Δ*smc*Δ*parAB*$p_{nat}$*parAB-HT*)**

The strains expressing *parAB-HT* under the control of the native promoter were constructed using pKP05 (pSS170 $p_{nat}$*parAB-HT*) integrative plasmid. To construct the pKP05 plasmid, the sequence encompassing the *parAB* operon with its promoter (2587 bp) was amplified using KP_66Fw/KP_43Rv primers and cosmid Sv-4-A09 as a template. The product was cloned by SLIC to the pSS170-*halotag* vector digested with XmaJI. The reaction mixture was used to transform *E. coli* DH5α cells, hygromycin hygromycin-resistant transformants were selected. The obtained vector pKP05 (pSS170 $p_{nat}$*parAB-HT*) was verified by PCR (using pSSseq_Fw/pSSseq_Rv primers), followed by sequencing the resulting product. The pKP05 vector was introduced to the wild strain S*. venezuelae* (NRRL B-65442), MD030 (Δ*parAB*) and KP004 (Δ*smc*Δ*parAB*) by intergeneric conjugation with *E. coli* ET12567/pUZ8002 followed by a selection of hygromycin-resistant exconjugates. The obtained strains KP005 (WT-$p_{nat}$*parAB*-HT), KP006 (Δ*parAB*$p_{nat}$*parAB*-HT) and KP007 (*Δsmc, ΔparAB,* $p_{nat}$*parAB*-HT) were verified by PCR (pSSseq_Fw/pSSseq_Rv primers) and by Western blotting using anti-HaloTag and anti-ParB antibody.

**Construction of *S. venezuelae* strains producing ParB-HT under the control of $p_{rtet}$ promoter: KP008 (WT $p_{rtet}$*parB-HT*), KP009 (Δ*parB* $p_{rtet}$*parB*-HT), KP010 (Δ*parB::apra* $p_{rtet}$*parB*-HT,) and KP011 (Δ*parB* $p_{rtet}$*parB*-HT*, ftsZ-ypet*)**

The strains expressing *parAB-HT* under the control of the $p_{rtet}$ promoter ($p_{tcp}$830 controlled by the reverse TetR repressor ($TetR_{rv}$) (Klotzsche et al., 2009)) were prepared using pKP08 (pSS170 $p_{rtet}$*parB-HT*). To construct pKP08, the sequence encoding the *tetRrv* gene (1068 bp) was amplified using KP_37Fw/KP_37BHTRv primers and pKP03 (pSS170 $p_{rtet}$*halotag*) plasmid as the template. At the same time, *parB* gene (1165 bp) was amplified using KP_43BHTFw/KP_43Rv primers and the Sv-4-A09 cosmid as a template. Both products were cloned to a vector digested with the restriction enzyme XmaJI using the Gibson Assembly method. A reaction mixture was used to transform *E. coli* DH5α and hygromycin-resistant transformants were selected. The obtained pKP08 (pSS170 $p_{rtet}$*parB-HT*) vector was verified by PCR reaction using pSSseq_Fw/pSSseq_Rv primers followed by sequencing of PCR product. The pKP08 vector was then introduced to the S . *venezuelae* wild type strain (NRRL B-65442), MD020 (Δ*parB*), MD002 (Δ*parB::apra*) by intergeneric conjugation with *E. coli* ET12567/pUZ8002 followed by selecting hygromycin-resistant exconjugates. The obtained KP008 (WT $p_{rtet}$*parB-HT*) KP009 (Δ*parB*, $p_{rtet}$*parB*-HT) and KP010 (Δ*parB*::*apra* $p_{rtet}$*parB-HT*) strains were verified by PCR using pSSseq_Fw/pSSseq_Rv primers. To obtain the KP011 strain (Δ*parB*, $p_{rtet}$*parB*-HT *ftsZ-ypet*), the pKF351 was introduced to KP009 by intergeneric conjugation followed by a selection of apramycin-resistant exconjugates.

**Construction of pBSK plasmid containing *parS* and mutated *parS* site**

To construct a pBSK plasmid containing a single *parS* site the fragment of the parAB promoter region (365 bp) from S. coelicolor was amplified with primers parA_fw and parAB_rv, using chromosomal S. coelicolor DNA as the template. The obtained PCR product was digested with HindIII and BamHI and cloned into the pBSK vector digested with the same enzymes. The obtained pBSK*parS* vector was verified by digestion and sequencing. To introduce mutation in the *parS* site, PCR was performed using the following two pairs of primers: parA_fw with parSA_mut rv and parSA_mut fw with parAB_rv and pBSK*parS* as a template. Next, the obtained products were mixed to serve as the template for overlap PCR with primers parA_fw and parAB_rv. The obtained PCR product was digested with HindIII and BamHI and cloned into the pBSK vector, digested with the same enzymes. The obtained pBSK$parS_{mut}$ vector was verified by digestion and sequencing.

**Quantification of the *oriC/arm* ratio**

To estimate the *oriC/*arm ratio, chromosomal DNA was extracted from *S. venezuelae* 5 ml cultures growing for 13-26 h or from 50 ml culture in MYM growing for 8 to 16 h. One ml of each culture was centrifuged (1 min at 5000 rpm and room temperature). The supernatant was discarded, and the mycelium was used for the subsequent isolation of chromosomal DNA using a Genomic Mini AX *Streptomyces* kit (A&A Biotechnology) according to the manufacturer's protocol. The purified DNA was dissolved in 50 µl of DNase-free water (Invitrogen). The DNA concentration was measured at 260 nm and subsequently diluted to a final concentration of 1 ng/ml.

qPCR was performed using Power Up SYBR Green Master Mix (Applied Biosystems) with 2 ng of chromosomal DNA serving as a template for the reaction and oligonucleotides complemented to the *oriC* region (*oriC*) (gyr2_Fd, gyr2_Rv) or the right arm termini (*arm*) (arg3_Fd, arg3_Rv) (Table S3). The changes in the *oriC/*arm ratio were calculated using the comparative ΔΔCt method, with the arm region being set as the endogenous control. The *oriC*/arm ratio was estimated as 1 in the culture (5 ml) growing for 26 h, corresponding to the appearance of spore chains.

**Western blotting**

For Western blotting analyses, *S. venezuelae* cultures were set up as follows: 5 ml of liquid MYM was inoculated with 1 ml of spores with $OD_{600}$ = 0.05, and the cultures were incubated for 2-26 h at 30°C with shaking (180 rpm). Next, a 2 ml sample of the culture was collected by centrifugation (5000 g, 5 min, 4°C), washed twice with phosphate-buffered saline (PBS buffer), resuspended in 200 µl of chilled PBS buffer supplemented with Pierce™ Protease Inhibitor Tablets (Thermo Fisher Scientific, USA), and disrupted by sonication. The cell lysate was then clarified by centrifugation (12 000 g, 5 min, 4°C), and the supernatant was transferred to a fresh tube. The total protein concentration was quantified using the ROTI®Quant Universal kit (Carl Roth, Germany). Cell lysate containing 10–20 µg of total protein was mixed with 6x SB buffer (375 mM Tris–HCl pH 6.8, 12% SDS, 0.06% bromophenol blue, 600 mM DTT, 60% glycerol), denatured at 95◦C for 10 min, and resolved by standard Laemmli acrylamide gel electrophoresis (SDS-PAGE). After electrophoresis, proteins were stained with InstantBlue Coomassie Protein Stain (Abcam, UK) (CBB, loading control) or transferred to a nitrocellulose membrane (Amersham, UK) and blocked with 2% skimmed milk (SM Gostyn,´ Poland) in Tris-buffered saline (TBS buffer) supplemented with 0.05% Tween-20 (TBST buffer). The blocked membrane was subsequently incubated with primary rabbit polyclonal antiparB antibody for ParB detection, mouse monoclonal anti-HaloTag antibody (Promega, US) for HaloTag detection or mouse monoclonal anti-FLAG antibody (Sigma−Aldrich, US) for FLAG-Tag detection. Primary antibodies were diluted 1:100 in TBST buffer, and the membrane was incubated for 1 hour at room temperature followed by washing three times with TBST buffer. Next, the membrane was incubated with secondary polyclonal antibodies: anti-mouse IgG conjugated with horseradish peroxidase (HRP, Invitrogen, US), anti-rabbit or anti-mouse IgG conjugated with horseradish peroxidase (Invitrogen, US) . The secondary antibodies were diluted 1:5000 (HRP-conjugated) in TBST buffer. Next, the membrane was washed as described earlier. For HRP signal detection, the membrane was washed briefly with 5 ml of SuperSignal West Pico PLUS Chemiluminescent Substrate solution (Thermo Fisher Scientific, USA), and the chemiluminescence signal was quantified using ChemiDoc XRS+ (Bio-Rad, USA). The band intensities were quantified using Fiji software.
